# GABA_B_ receptors mediate presynaptic inhibition of proprioceptive afferents and have reduced action after spinal cord injury

**DOI:** 10.64898/2026.01.21.700955

**Authors:** K. Metz, K. Hari, A. Lucas-Osma, R. Mangukia, TO. Ayantayo, I. Concha-Matos, Y. Sun, JF. Yang, DJ. Bennett, MA. Gorassini

## Abstract

Despite a long history of studying presynaptic inhibition of the Ia afferent synapse that produces the monosynaptic EPSP on motoneurons, recent evidence has upset the conventional idea that GABA_A_ receptors mediate this inhibition and instead suggests that there are mainly GABA_B_ receptors at this synapse. However, without targeted access to the GABAergic neurons that activate these receptors, quantifying their functional contribution to presynaptic inhibition has proven difficult. We demonstrate here that focal optogenetic activation of terminals of a subpopulation of GAD2+ GABAergic neurons that exclusively project ventrally to Ia afferent synapses produce long-lasting presynaptic inhibition that is entirely mediated by GABA_B_ receptors and simultaneously produces a characteristic brief GABA_A_ receptor-mediated IPSP on the motoneurons. These ventral GAD2 neurons are recurrently activated by Ia afferents, contributing to post-activation depression with repeated afferent reflex testing, with a similar long time-course to post-activation depression of the H-reflex induced in humans from either repetitive activation of the same Ia afferents or from antagonist nerve conditioning. In contrast, focal activation of dorsally projecting GAD2 neurons does not directly cause presynaptic inhibition or postsynaptic IPSPs but does produce primary afferent depolarization. Following chronic spinal cord injury (SCI), the expression of GABA_B_ receptors on the Ia terminal is halved, and in mice and humans, is associated with a similar decrease of GABA_B_ receptor-mediated post-activation depression of Ia-EPSPs transmission, which is reversed by the GABA_B_ receptor agonist baclofen. In summary, GABA_B_ receptors mediate presynaptic inhibition, but are down regulated with SCI, contributing to reflex hyperexcitability associated with spasticity.

**Key Points Summary:**

- Presynaptic inhibition of Ia afferents is mediated by the recurrent activation of terminal GABA_B_ receptors by a subpopulation of ventrally projecting GAD2+ interneurons.
- In contrast, dorsally projecting GAD2+ interneurons activate GABA_A_ receptors on Ia afferent nodes to facilitate action potential conduction through branchpoints.
- Repetitive activation of Ia afferents at rates of every 10 s or faster produces post-activation depression via neurotransmitter depletion and from activation of terminal GABA_B_ receptors.
- These ventrally projecting GAD2+ interneurons can also be activated by other afferents that then produce PAD-evoked spikes to produce post-activation depression from conditioning nerve stimulation.
- The reduction of GABA_B_ receptors on the Ia terminal in spinal cord injury results in reduced presynaptic inhibition and post-activation depression, contributing to reflex hyperexcitability.

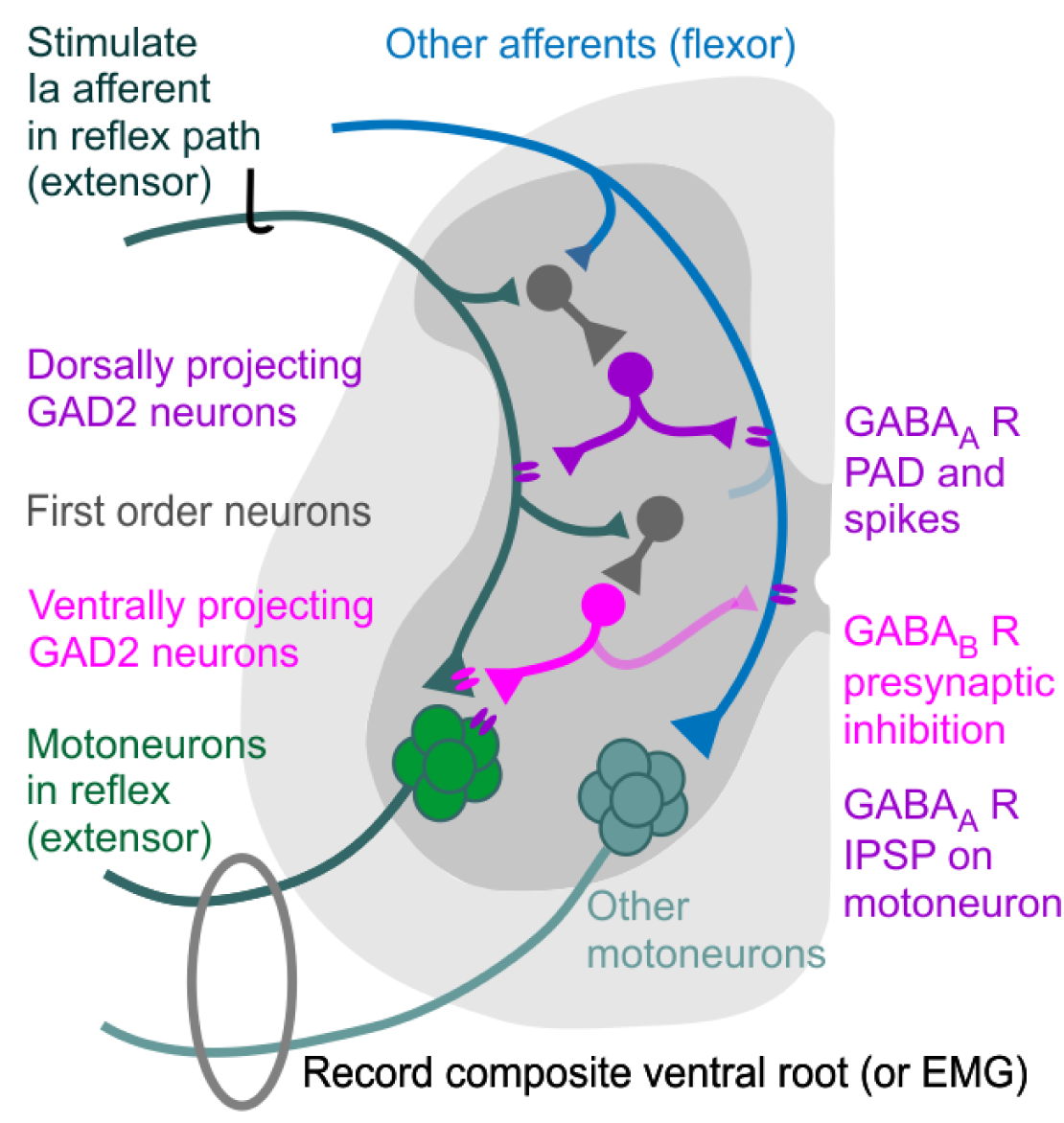
Abstract legend: Schematic of GABAergic circuit producing presynaptic inhibition and primary afferent depolarization (PAD) in proprioceptive Ia afferents. We propose two populations of GAD2+ GABAergic interneurons, one with dorsal projections (purple) that activate GABA_A_ receptors on the nodes of Ia afferents to produce PAD and subsequent facilitation of Ia afferent conduction, and another ventrally projecting population (pink) that activates GABA_B_ receptors on the Ia afferent terminal to produce presynaptic inhibition via inhibition of VCa^2+^ channels and reduction of neurotransmitter release and replenishment. Both are activated by first order interneurons (grey). Repetitive activation of Ia afferents (green extensor) recurrently activates the ventrally projecting GAD2+ neurons to activate terminal GABA_B_ receptors and long-lasting post-activation depression of Ia EPSPs and reflexes as measured from ventral root recordings. Strong conditioning stimulation of other afferents (blue flexor) activates dorsal GAD2+ neurons that can produce PAD-evoked spikes in extensor afferents that orthodromically activate motoneurons to set up post-activation depression of subsequent extensor reflexes. Here, PAD is also evoked in other afferents (flexor) by dorsally projecting GAD2+ neurons (light pink branch) but without activation of the ventrally projecting GAD2+ neurons or presynaptic inhibition.

## Introduction

After injury to the spinal cord (SCI), abnormal control over sensory transmission to the brain and spinal cord can lead to a host of devastating secondary complications like neuropathic pain (Park *et al*., 2018) and spasticity (Rabchevsky & Kitzman, 2011; Silva *et al*., 2014).

Excessive transmission from proprioceptive Ia afferents to spinal interneurons and motoneurons may contribute to the generation of muscle spasms, hyperreflexia, clonus and hypertonia in spasticity [reviewed in (Lalonde & Bui, 2021)]. One mechanism that has been proposed to control the transmission of Ia afferents to their target motoneurons is presynaptic inhibition, broadly defined as a reduction of neurotransmitter release in a central synapse (Quevedo, 2009). However, despite its nearly one hundred-year-old history, presynaptic inhibition has not been directly measured in Ia afferent terminals, unlike in other axons where presynaptic terminals and associated transmitter release mechanisms have been directly studied with dual patch clamp recordings (Sakaba & Neher, 2001b), leaving much uncertainty as to whether it occurs and what receptors mediate it [reviewed in Supplemental Material of (Hari *et al*., 2022)]. While Ia afferent terminals are extensively innervated by GAD2+ GABAergic neurons (Betley *et al*., 2009), recent evidence has shown that these terminals lack functional GABA_A_ receptors that were widely believed to causes presynaptic inhibition. Thus, the first goal of this paper is to employ focal optogenetic release of GABA from GAD2+ neurons onto Ia afferents to examine the receptors involved in presynaptic inhibition of Ia synaptic transmission to motoneurons. We describe here a distinct subpopulation of ventral GAD2 neurons that innervate Ia afferent terminals and produce presynaptic inhibition by terminal GABA_B_, rather than GABA_A_, receptors (Fig. 1A). In addition, we found that more dorsally located GAD2 neuron populations mediate GABA_A_ receptor actions on Ia afferents that are readily mistaken for presynaptic inhibition during repetitive or conditioning stimulation protocols commonly used to assess presynaptic inhibition, which we also clarify.

**Figure 1:**
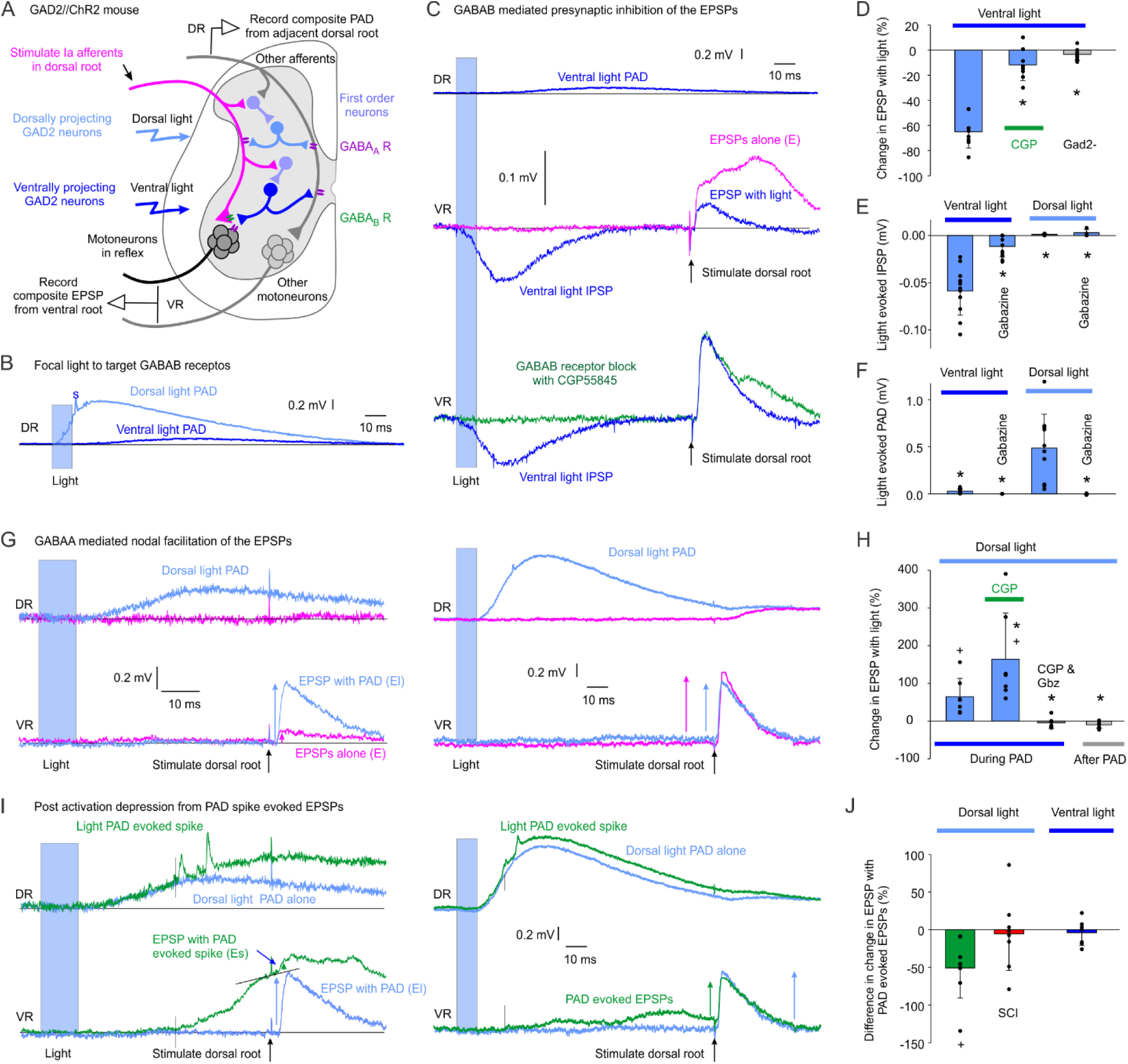
**A)** Schematic of dorsal root (DR) and ventral root (VR) recordings in response to low-intensity stimulation of Ia afferents in adjacent DR (pink) and optogenetic activation of dorsal (light blue) and ventral (dark blue) projecting GAD2 neurons by blue light in GAD2//ChR2 mice. Dorsal GAD2 neurons innervate GABA_A_ receptors (purple) near Ia nodes to evoke PAD and ventral GAD2 neurons innervate GABA_B_ receptors (green) at Ia terminal to produce presynaptic inhibition, and GABA_A_ receptors on motoneuron to produce an IPSP. Excitatory interneurons between Ia afferent and GAD2 neurons are coloured in lavender. **B)** DR response to a focused 10 ms light pulse [447nm laser, 0.7 mW/mm^2^, 1.5x light threshold (LT) to evoke PAD or motoneuron responses] applied to dorsal and ventral location of the whole sacral spinal cord (“s” indicates PAD-evoked spike). **C)** Top: DR response to the same ventral light pulse alone.

Repetitive activation of the Ia afferent terminal is thought to induce presynaptic inhibition (Eccles & Rall, 1951), as indirectly assessed by a reduction of the monosynaptic, Ia-mediated excitatory postsynaptic potential (EPSP) in a motoneuron that can last for up to 10 seconds (Lloyd & Wilson, 1957). The reduction of the Ia-EPSP by a prior activation of the same EPSP, with the first and subsequent EPSPs evoked by the same Ia afferent population each time, has been referred to as homosynaptic, post-activation depression (Crone & Nielsen, 1989; Hultborn *et al*., 1996). The magnitude of post-activation depression is dependent upon the rate of repetitive stimulation of the Ia-EPSP synapse, thereby termed rate-dependent depression (RDD), where the suppression of Ia-EPSPs or Ia-mediated monosynaptic reflexes increases as the stimulation frequency increases (Lloyd & Wilson, 1957; Ishikawa *et al*., 1966; Jiang *et al*., 2015). In humans, it is thought that post-activation depression of Ia-EPSPs is suppressed after SCI, as reflected by a reduced suppression of H- or posterior root reflexes in response to repetitive activation of Ia afferents (Ashby & Verrier, 1975; Calancie *et al*., 1993; Nielsen *et al*., 1993; Schindler-Ivens & Shields, 2000; Grey *et al*., 2008; Kumru *et al*., 2015; Hofstoetter *et al*., 2019).

The effect of repetitively activating a central synapse on neurotransmitter vesicle release and its replenishment rate during post-activation depression has been mainly studied in the Calyx of Held synapse due to the large size of the presynaptic terminal (Neher & Sakaba, 2001; Sakaba & Neher, 2001b). By voltage clamping the presynaptic terminal to regulate the amount calcium entry and vesicle release, while simultaneously measuring the EPSCs in the postsynaptic target neuron, it was estimated that full recovery of the readily releasable pool of neurotransmitters (vesicle replenishment) occurs when repetitive activation of the synapse is separated by at least 10 seconds (Sakaba & Neher, 2001a). Remarkably, the profile of vesicle replenishment closely matches the profile of Ia-EPSP and H-reflex recovery during RDD, whereby low levels of vesicle replenishment (and greater RDD) occur at stimulation rates faster than 1 second, followed by a more gradual increase to full vesicle replenishment (and no RDD) by 10 seconds. Moreover, the amount of vesicle replenishment is reduced when the levels of calcium or calmodulin were lowered in the presynaptic terminal (Sakaba & Neher, 2001a). A lowering of vesicle replenishment could increase presynaptic inhibition and its effect on post-activation depression, especially at fast repetition rates, as there would be less neurotransmitter vesicles available to evoke Ia-EPSPs following each stimulation. As described below, factors that affect calcium entry into the presynaptic terminal, and its activation of calmodulin, could affect both vesicle replenishment and post-activation depression of the Ia-EPSP.

Recently, we have demonstrated that GABA_B_ receptors are mainly localized at the presynaptic terminal of the Ia afferent (Hari *et al*., 2022). Because GABA_B_ receptors inhibit voltage-activated calcium channels (VCa^2+^) in the presynaptic terminal (Dolphin & Scott, 1986; Lev-Tov *et al*., 1988; Curtis *et al*., 1997; Abe *et al*., 2019), and likely the activation of Ca^2+^-calmodulin, their activation might reduce vesicle release and replenishment, as shown in the Calyx of Held (Sakaba & Neher, 2003). Thus, the activation of GABA_B_ receptors should increase presynaptic inhibition and post-activation depression of Ia-EPSPs during fast repetitive stimulation. Based on recent anatomical data (Lin *et al*., 2023), we have proposed that the activation of GABA_B_ receptors on the Ia terminal during its repetitive stimulation could occur via collaterals of the Ia afferent that synchronously activate a recurrent inhibitory circuit containing GABAergic (GAD2+) neurons that in turn project onto the terminals of the same Ia afferent [Fig. 1A, see also (Metz *et al*., 2023a)], likely through a trisynaptic circuit involving Ia activation of first order neurons that widely innervate the GAD2 neurons. This recurrent inhibitory pathway could activate GABA_B_ receptors on the Ia terminal and facilitate presynaptic inhibition via inhibition of VCa^2+^ channels. Thus, the second goal of this paper is to explore the role of GABA_B_ receptors in mediating presynaptic inhibition produced by *repetitive* Ia afferent stimulation. While we show here that repeated focal optogenetic activation of GAD2 neuron terminals innervating Ia afferents causes a profound inhibition of the Ia-EPSP that is entirely mediated by GABA_B_ receptors, repeated activation of the Ia afferents themselves produces an even larger inhibition that is only partly mediated by GABA_B_ receptors, and likely involves direct neurotransmitter depletion from the Ia afferent terminal, similar to that shown in the Calyx of Held (Sakaba & Neher, 2001b).

Post-activation depression of Ia-EPSPs is reduced after SCI as described above, prompting the question of whether GABA_B_ receptor action on Ia afferent terminals is also reduced. Indeed, the first line of treatment for hyperreflexia and spasms after SCI is the GABA_B_ receptor agonist baclofen that potently suppresses Ia-mediated monosynaptic reflexes (Dietz *et al*., 2023). Furthermore, when activating GABA_B_ receptors with baclofen, the monosynaptic reflex is reduced less in rodents with chronic SCI compared to in controls (Li *et al*., 2004b), likely via impaired GABA_B_ receptor actions including inhibition of terminal VCa^2+^ channels and vesicle release (Dolphin & Scott, 1986; Lev-Tov *et al*., 1988; Curtis *et al*., 1997; Abe *et al*., 2019). Thus, the third goal of this paper is to examine how GABA_B_ receptor actions on Ia afferents change with SCI in mice and humans. By employing confocal imaging, optogenetic GAD2 activation, direct afferent recordings and pharmacological receptor blockers or agonists in mice, we examine whether GABA_B_ receptors on Ia afferent terminals are down regulated with SCI and whether this contributes to the reduction in post-activation depression and RDD of Ia-EPSPs with SCI. We also explore whether a similar reduced GABA_B_ receptor action occurs in humans with SCI by examining changes in post-activation depression (and RDD) of H-reflexes during facilitation of GABA_B_ receptors with baclofen.

In addition to repetitively activating the same Ia-motoneuron synapse, the suppression of Ia-EPSPs is also produced by a prior conditioning stimulation to a separate nerve that is likely also produced by presynaptic inhibition of the Ia terminal (Eccles *et al*., 1961a; Willis, 2006). Originally, this form of presynaptic inhibition was thought to be mediated by the depolarization of the Ia terminal (primary afferent depolarization or PAD) by GABA_A_ receptors that either shunted terminal action potentials or inactivated sodium channels to reduce neurotransmitter release (Eccles, 1964; Rudomin & Schmidt, 1999). However, as described above, the finding that there is very little GABA_A_ receptors and PAD at the Ia terminal required a different mechanism to explain the Ia-EPSP suppression by the conditioning nerve stimulation (Lucas-Osma *et al*., 2018; Hari *et al*., 2022). We and others have shown that stimulation of afferents in one dorsal root can produce PAD in the nodes of Ia afferents in another dorsal root that, if large enough, can evoke orthodromic spikes which travel to the Ia terminal and activate a monosynaptic Ia-EPSP in the motoneuron (Eccles *et al*., 1961b; Duchen, 1986; Willis, 1999; Hari *et al*., 2022; Metz *et al*., 2023a). Thus, like direct repetitive activation of the Ia afferent, a conditioning nerve stimulation may also produce presynaptic inhibition and post-activation depression given that the same Ia-EPSP synapse is repetitively activated by the conditioning, and then test, afferent stimulation. To examine if suppression of Ia-EPSPs by a conditioning nerve stimulation is also reduced after SCI in a manner similar to repetitive activation of the same Ia afferents, we examined here if the suppression of soleus (extensor) H-reflexes by a prior common peroneal nerve (CPN, flexor) stimulation was reduced compared to uninjured control participants (El-Tohamy & Sedgwick, 1983; Roby-Brami & Bussel, 1990; Metz *et al*., 2023a).

Collectively, our results suggest a role of GABA_B_ receptor activation in facilitating presynaptic inhibition of the Ia afferent terminal that is reduced after SCI due, in part, to a reduction of GABA_B_ receptors activated by a distinct population of ventral GAD2 neurons. This reduced activation of GABA_B_ receptors on the Ia afferent terminal reduces presynaptic inhibition and post-activation depression (RDD) of the Ia-motoneuron synapse, likely contributing to hyperreflexia in spasticity.

## Methods

### Animals and ethics

Recordings were made from large proprioceptive group Ia sensory afferents and motoneurons in adult mice (2.5–6 months old, both female and male equally). Animals were caged in groups of 2-4 and maintained on a 12-hour light/dark cycle, in stable conditions of temperature and humidity, with food and water ad libitum. All experimental procedures were approved by the University of Alberta Animal Care and Use Committee, Health Sciences division (ACUC protocol nos. AUP00000224 and AUP00002891) in accordance with the Canadian Council on Animal Care guidelines. We evaluated GABAergic neurons in a strain of mice with Cre expressed under the endogenous Gad2 promotor region. Gad2 encodes the Glutamate decarboxylase 2 enzyme GAD2 (also called GAD65), which is unique to axoaxonic contacting GABAergic neurons that project to afferents, whereas all GABAergic neurons express GAD1 (Betley *et al*., 2009). These GAD2+ neurons were activated or inhibited optogenetically using channelrhodopsin-2 (ChR2) (Zhang *et al*., 2011; Pinol *et al*., 2012) or archaerhodopsin-3 (Ach3) (Chow *et al*., 2010; Kralj *et al*., 2011), respectively. The following mouse strains were employed:

1. *Gad2^tm1(cre/ERT2)Zjh^* mice (abbreviated Gad2^CreER^ mice; The Jackson Laboratory, Stock # 010702; CreER^T2^ fusion protein expressed under control of the endogenous Gad2 promotor) (Taniguchi *et al*., 2011),
2. *B6;129S-Gt(ROSA)26Sort^m32(CAG-COP4*H134R/EYFP)Hze^* mice (abbreviated R26^LSL-ChR2-EYFP^ mice; The Jackson Laboratory, Stock # 012569; ChR2-EYFP fusion protein expressed under the R26::CAG promotor in cells that co-express Cre because a loxP-flanked STOP cassette, LSL, prevents transcription of the downstream ChR2-EYFP gene) (Madisen *et al*., 2012), and
3. *B6;129S-Gt(ROSA)26Sor^tm35.1(CAG-aop3/GFP)Hze^*mice (abbreviated R26^LSL-Arch3-GFP^ mice; The Jackson Laboratory Stock # 012735; Arch3-GFP fusion protein expressed under the R26::CAG promotor in cells that co-express Cre) (Madisen *et al*., 2012).

Heterozygous GAD2^CreER^ mice (i.e., GAD2^CreER/+^ mice) were crossed with homozygous reporter strains to generate GAD2^CreER/+^; R26^LSL-ChR2-EYFP^, and GAD2^CreER/+^; R26^LSL-Arch3-GFP^ mice that we abbreviate: GAD2//ChR2, GAD2//ChR2-EYFP, and GAD2//Arch3 mice. Offspring without the GAD2^CreER^ mutation, but with the effectors ChR2, or Arch3 were used as controls.

For all mice, genotyping was performed according to the Jackson Laboratories protocols by PCR of ear biopsies using primers specific for the appropriate mutant and wild type alleles for each of the mouse lines (see Supplementary Table 2 in Hari et al, 2022 for primer details).

CreER is an inducible form of Cre that requires tamoxifen to activate (Feil *et al*., 1997), which we applied in adult mice to prevent developmental issues of earlier induction of Cre.

Specifically, mice were injected at 4 - 6 weeks old with two doses of tamoxifen separated by two days, and studied > 1 month later, long after washout of tamoxifen. Each injection was 0.2 mg/g wt (i.p.) of tamoxifen dissolved in a corn oil delivery vehicle (Sigma C8267).

Mice were also studied following chronic spinal cord injury (SCI) where the S2 sacral spinal cord was transected at ∼2 months of age. Briefly, under general anesthetic (ketamine 90mg/kg and xylazine 12mg/kg i.p., and isoflurane) and sterile conditions, a laminectomy was performed on the L2 vertebrae to expose the S2 spinal cord. The dura was slit transversely, and 0.02 - 0.03 ml of xylocaine (2%) was applied topically. Using a surgical microscope, the spinal cord was transected by first making a small incision of the cord lateral to the dorsal vein, and while holding the pia with forceps spinal cord tissue was removed using suction (with a 1-cc syringe that had been melted and pulled to a fine tip). Caution was taken to avoid damaging the anterior artery or posterior/dorsal vein, because the sacrocaudal spinal cord dies without this midline vasculature. The dura was closed with a single 8-0 silk suture, the muscle layers and skin were tightly sutured over the cord and the mouse was allowed to recover. Buprenorphine (0.063 mg/kg s.c.) was administered immediately after surgery and in the morning on the following 2 days. All experiments were performed 2–4 months after injury, when tail-muscle spasticity was fully developed, and mice were considered to be in a chronic spinal state [*chronic spinal state,* (Bennett *et al*., 1999)]. Non-spastic mice with inactive tails were excluded. Chronic spinal mice were compared to age-matched, uninjured mice (mice were randomly assigned to these two groups).

### Animal experimental procedures

#### Ex vivo recordings from axons and motoneurons in whole adult spinal cords

Mice were anaesthetized with urethane (0.11 g per 100 g, with a maximum dose of 0.065 g); a laminectomy was performed and then the entire sacrocaudal spinal cord was rapidly removed to a dissection chamber and immersed in oxygenated modified artificial cerebrospinal fluid (mACSF) (Hari *et al*., 2022). The mACSF was composed of (in mM) 118 NaCl, 24 NaHCO3, 1.5 CaCl2, 3 KCl, 5 MgCl2, 1.4 NaH2PO4, 1.3 MgSO4, 25 D-glucose and 1 kynurenic acid. Spinal roots were removed, except the sacral S3, S4 and caudal Ca1 ventral roots (VRs) and dorsal roots (DRs) on both sides of the cord. After 1.5 hours in the dissection chamber (at 20°C), the cord was transferred to a recording chamber containing normal ACSF (nACSF) composed of (in mM) 122 NaCl, 24 NaHCO3, 2.5 CaCl2, 3 KCl, 1 MgCl2 and 12 D-glucose, and maintained at 23–32 °C, with a flow rate >3mlmin−1. Both the mACSF and nACSF were saturated with 95% O2/5% CO2 and maintained at pH 7.4. A 1-hour period in nACSF was given to wash out the residual anaesthetic before recording, at which time the nACSF was recycled in a closed system. The cord was secured onto tissue paper at the bottom of a rubber (SilGuard) chamber by insect pins placed into connective tissue and cut root fragments. The spinal cord was oriented with its left side upwards when applying dorsal and ventral light and making recordings from afferents in the DR and motoneurons in the VR. This preparation is particularly useful as the small sacrocaudal spinal cord is the only portion of the adult spinal cord that survives whole ex vivo, with very similar spinal circuitry, reflex and motoneuron properties to those seen in the lumbar hindlimb region of other preparations, including Ia afferent innervation of muscle spindles and monosynaptic reflexes (Bennett *et al*., 1999; Bennett *et al*., 2001; Murray *et al*., 2010; Murray *et al*., 2011).

#### Ia monosynaptic EPSPs

The composite EPSPs in many motoneurons were recorded from the central cut end of ventral roots mounted in grease (grease gap), which yields reliable estimates of the motoneuron EPSPs, although attenuated by the distance from the motoneuron pool (Luscher *et al*., 1979; Hari *et al*., 2022). Freshly cut dorsal and ventral roots were mounted onto silver–silver chloride wires just above the bath and covered in grease over a ∼2-mm length. Return and ground wires were in the bath and likewise made of silver–silver chloride. The high impedance seal on the motor axons reduces extracellular currents, allowing the recording to reflect intracellular potentials (Luscher *et al*., 1979; Stein, 1980). To evoke the Ia monosynaptic EPSP, proprioceptive afferents in the DR were stimulated with a constant current stimulator using a 0.1ms pulse width at 1.1× T (ISO-Flex), where T is the EPSP threshold and is similar to the afferent volley threshold. To induce post-activation depression, the DR was stimulated four times (M1 to M4) at a rate of every 60 - 100 ms. The EPSPs were identified as monosynaptic by their rapid onset (first component, ∼1ms after afferent volley arrives in the ventral horn), lack of variability in latency (< 1-ms jitter), persistence at high rates (10 Hz) and appearance in isolation at the threshold (T) for evoking EPSPs with DR stimulation (< 1.1× T; T, afferent volley threshold), unlike polysynaptic reflexes, which vary in latency, disappear at high rates and mostly need stronger DR stimulation to activate. Post-activation depression was compared in control mice at the 60 - 100 ms stimulation rate during GAD2+ neuron silencing or activation, and during GABA_B_ receptor blockade (see next section below). To match the H-reflex data in human participants, the rate-dependency of post-activation depression (RDD) was assessed at a stimulation rates every 0.5, 1, 2 and 5 s and compared between control and chronically injured mice with and without GABA_B_ receptor blockade with CGP55845.

#### Activation and inactivation of GAD2+ neurons and GABA_A_ and GABA_B_ receptor modulation

The GAD2//Arch3 mice were used to optogenetically inhibit GAD2+ neurons during post-activation depression (with a 532 nm LRS-0532-GFM-00200-01 laser from Laserglow Technologies) and GAD2//ChR2 mice were used to excite the GAD2+ neurons (with a 447 nm D442001FX laser). Light was derived from the laser beam passed through a fiber optic cable (MFP_200/240/3000-0.22_2m_FC-FC, Doric Lenses) and then a half cylindrical prism the length of about two spinal segments (8mm; 3.9 mm focal length, Thor Labs), which collimated the light into a narrow long beam (200 µm wide and 8 mm long). This narrow beam was focused longitudinally on the left side of the spinal cord. This beam was either focused on the dorsal horn, to target the epicenter of GAD2 neurons, which are dorsally located (Hari *et al*., 2022), or on the lamina IX ventral horn to target GAD2 terminals over the motoneurons. Ventral GAD2+ neuron terminals were selectively activated in GAD2//ChR2 mice by applying blue light to the ventral surface of the spinal cord in this way, using 10 ms light pulses applied every 100 ms either with or without DR stimulation. Three pulses were applied before the test Ia-EPSP. Arch3 is a proton pump that is activated by green light, leading to a hyperpolarization and slowly increased pH (over seconds), both of which inhibit the neurons (Zhang *et al*., 2011; El-Gaby *et al*., 2016). Thus, long light pulses (∼800ms) were used to inhibit GAD2+ neurons in GAD2//Arch3 mice during post-activation depression. CGP55845 (abbreviated CGP in figures), a GABA_B_ receptor antagonist (0.3 µM), gabazine, a GABA_A_ receptor antagonist (50 µM) and baclofen, a GABA_B_ receptor agonist (1 μM) (all from Tocris), were added to the nACSF to examine their effects on post-activation depression and Ia hyperpolarization. Drugs were first dissolved as a 10–50mM stock in water or DMSO before final dilution in nACSF. DMSO was necessary for dissolving CGP55845, but it was kept at a minimum (final DMSO concentration in ACSF < 0.04%), which by itself had no effect on reflexes or sensory axons in vehicle controls (Hari *et al*., 2022).

#### DC recordings of dorsal roots during baclofen application

The hyperpolarizing effect of the GABA_B_ receptor agonist baclofen was measured from the cut end of the DR. As with VR recordings, the same grease gap method was used to record the DC membrane potential of many sensory axons in the DR, including the large Ia afferent that dominate the signal (Hari *et al*., 2022). The DR recordings were amplified (2,000 times), high-pass filtered at 0.1Hz to remove drift, low-pass filtered at 10 kHz and sampled at 30 kHz (AxoScope 8; Axon Instruments/ Molecular Devices). Recordings were made during blockade of fast synaptic transmission to synaptically isolate the axons before adding baclofen. Data from 6 DRs in two mice were used in each group.

#### Data analysis

To quantify the suppression of Ia EPSPs during repetitive stimulation of the DR and/or from repetitive ventral GAD2+ activation, the amplitude of the fourth EPSP (M4) in the stimulus train was compared to the first EPSP (M1) by the formula % change EPSP = 100% x (M4 – M1) / M1). In control and SCI mice the hyperpolarizing effect of the GABA_B_ receptor agonist, baclofen, on the mean membrane potential of the Ia afferents was measured as the decrease in DC membrane potential from baseline measured 20 minutes after the application of baclofen (peak). Data from different DRs was used as independent measures in each mouse.

#### Viral labeling of sensory afferents

Large-diameter peripheral afferents were labeled by viral injections (Foust *et al*., 2008; Li *et al*., 2020; Hari *et al*., 2022) to quantify the number of GABA_B_ receptors on Ia afferent terminals. Adeno-associated viral vectors (AAVs) with the transgene encoding the cytoplasmic fluorophore tdTom under the CAG promoter were injected i.p. into anesthetized P1–2 mice (AAV9-CAG-tdTom, 5.9×1012 viral genomes per milliliter (vg per ml); 2–4µl per injection; UNC Vector Core). Mice were perfused for immunolabeling > 60 days after injection (adult mice). This injection method yields labeling of afferents in each spinal segment, without central labeling of other neurons (with the exception of one or two motoneurons labeled per spinal segment), allowing afferents to be traced to the motoneurons for morphological identification as Ia afferents.

### GABA_B_ receptor quantification on Ia afferent terminals from immunohistochemistry

#### Tissue fixation and sectioning

GAD2//ChR2-EYFP mice whose afferents were labeled genetically by a previous AAV9-CAG-tdTom viral i.p. injection were euthanized with Euthanyl (Bimeda-MTC; 700mg kg−1) and perfused intracardially with 10ml of saline for 3–4 minutes, followed by 40ml of 4% PFA (in 0.1M phosphate buffer at room temperature) over 15 minutes (Gabra5-KO mice also fixed similarly). The spinal cords of these mice were post-fixed in PFA for 1hour at 4 °C and then cryoprotected in 30% sucrose in phosphate buffer (∼ 48hours). After cryoprotection, all cords were embedded in OCT (Sakura Finetek), frozen at −60 °C with 2-methylbutane, cut on a cryostat CM3050 S (Leica Biosystems) in transverse 25-µm sections and mounted on slides. Slides were frozen until further use.

#### Immunolabeling

The tissue sections on slides were first rinsed with PBS (100mM, 10min) and then again with PBS containing 0.3% Triton X-100 (PBS-TX, 10-minute rinses used for all PBS-TX rinses). For all tissue, non-specific binding was blocked with a 1-hour incubation in PBS-TX with 10% normal goat serum (NGS; S-1000, Vector Laboratories) or normal donkey serum (NDS; ab7475, Abcam). Sections were then incubated for at least 20 hours at room temperature with the combination of the following primary antibodies in PBS-TX with 2% NGS or NDS: guinea pig anti-VGLUT1 (1:1,000; AB5905, Sigma-Aldrich), rabbit anti-GABAB1 receptor subunit (1:200; 322 102, Synaptic Systems), goat anti-RFP (1:500, orb334992, Biorbyt), and chicken anti-GFP (1:500, ab13970, Abcam), the latter two to amplify the tdTom and EYFP signals.

The next day, tissue was rinsed with PBS-TX with 2% NGS or NDS (3×10minutes) and incubated with fluorescent secondary antibodies. The secondary antibodies used included: donkey anti-rabbit Alexa Fluor 555 (1:500; ab150131, Abcam), donkey anti-rabbit Alexa Fluor 647 (1:500; ab150075, Abcam), donkey ant-chicken AF488 (1:500; 703 545 155, Jackson Immuno), goat anti-rabbit Pacific Orange (1:250; P31584, Thermo Fisher Scientific), goat anti-chicken Alexa Fluor 488 (1:500; ab96947, Abcam), goat anti-guinea pig Alexa Fluor 647 (1:500; A21450, Thermo Fisher Scientific), applied on slides for 2 hours at room temperature. After rinsing with PBS-TX (2×10minutes each) and PBS (2×10minutes each), the slides were covered with Fluoromount-G (00-4958-02, Thermo Fisher Scientific) and coverslips (no. 1.5, 0.175mm, 12-544-E, Thermo Fisher Scientific).

Standard negative controls in which the primary antibody was either (1) omitted or (2) blocked with its antigen (quenching) were used to confirm the selectivity of the antibody staining, and no specific staining was observed in these controls. Previous tests detailed by the manufacturer further demonstrate the antibody specificity, including quenching, immunoblots (western blots), co-immunoprecipitation and/or receptor knockout. Most antibodies had been previously verified for selectivity, as detailed in the manufacturer’s literature and other publications using knockout mice, as detailed in (Hari *et al*., 2022).

### Confocal and epifluorescence microscopy

Image acquisition was performed by confocal (Leica TCS SP8 confocal system) for high-magnification three-dimensional (3D) reconstruction. All the confocal images were taken with a ×63 (1.4 NA) oil immersion objective lens and optical sections were collected into a Z-stack over 10–20 µm. Excitation and recording wavelengths were set to optimize the selectivity of imaging the fluorescent secondary antibodies. The same parameters (laser intensity, gain and pinhole size) were used to take pictures for each animal, including the negative controls. Complete transversal sections were imaged with a ×20 (0.75 NA) objective lens using the Tile Scan option in the Leica Application Suite X (Leica Microsystems).

The fluorescently labeled afferents (tdTom), GAD2 positive cells (EYFP), GABA B1 receptors and VGLUT1 were analyzed using 3D reconstruction tools in Leica Application Suite X (Leica Microsystems) (Lucas-Osma *et al*., 2018). To minimize non-specific antibody staining, a threshold was set for each channel at a level corresponding to less than 10% background signal, determined from control sections processed without primary antibodies. Signals exceeding this threshold were rendered in 3D. GABA_B_ receptors located within the volume of the tdTomato-labelled Ia axon terminal (defined by a binary mask) were labeled in 3D reconstructions.

Receptor expression was quantified as mean fluorescence intensity within the reconstructed volumes. For comparisons between spinal cord–injured (SCI) and control mice, injured and control spinal cord sections were processed, mounted on the same slide, and imaged under identical acquisition settings. Mean GABA_B_ receptor intensity on Ia terminals in SCI mice was normalized to the mean intensity measured on Ia terminals in intact control mice and expressed as a percentage. Similarly, mean GABA_B_ receptor intensity on motoneurons in SCI mice was normalized to the corresponding intensity measured on motoneurons from control mice.

To assess the relative distribution of GABA_B_ receptors on the postsynaptic compartment, mean receptor intensity on Ia terminals was expressed as a percentage of the mean receptor intensity measured on motoneurons for both control and SCI mice. In addition, the proportion of Ia terminals expressing GABA_B_ receptors was calculated by dividing the number of VGLUT1-identified Ia terminals containing detectable GABA_B_ receptor signal by the total number of identified Ia terminals and expressed as a percentage.

### Human Participants and ethics

Experiments were approved by the Human Research Ethics Board at the University of Alberta (Protocols 000780557 and 00031413) and performed with informed consent of the participants. Our sample comprised of 36 participants with a SCI (26 male) ranging in age from 20 to 67 years [40.610 (15.743), Supplemental Table 1]. The age of injury varied from 0.5 to 33.5 years [8.117 (7.349)] with neurologic injury levels ranging from C2 to T12 with 16 of the 36 participants with SCI unable to voluntarily activate one or more muscles of the leg (Supplementary Table 1). Comparative data from 33 control participants (15 male) with no known neurological injury or disease were recruited. Ages of the control participants ranged from 20 to 58 years [29.817 (12.018)] and were significantly younger than the SCI group (P < 0.001, Mann-Whitney Rank Sum).

### Human experimental procedures

Participants were typically seated in their wheelchairs with one leg slightly extended to access the popliteal fossa. The right leg was used in all control and SCI participants except in 4 SCI participants where the more spastic left leg was used. Padded supports were added to facilitate complete relaxation and minimal movement of the leg. Participants were asked to rest completely with no talking and no hand or arm movements, including participants with motor complete SCI, as residual pathways may exist and movement of the upper body could stretch or shift the lower body, affecting H-reflex measurements.

#### EMG recordings

A pair of Ag-AgCl electrodes (Kendall; Chicopee, MA, USA, 3.2 cm by 2.2 cm) was used to record surface EMG from the soleus and tibialis anterior (TA) muscles with a ground electrode placed just below the knee. The EMG signals were amplified by 1000 and band-pass filtered from 10 to 1000 Hz (Octopus, Bortec Technologies; Calgary, AB, Canada) and then digitized at a rate of 5000 Hz using Axoscope 10 hardware and software (Digidata 1400 Series, Axon Instruments, Union City, CA).

#### Nerve stimulation to evoke an H-reflex

The tibial nerve (TN) was stimulated in a bipolar arrangement using a constant current stimulator (1 ms rectangular pulse width, Digitimer DS7A, Hertfordshire, UK) to evoke an H-reflex in the soleus muscle. After searching for the TN with a probe over the skin, an Ag-AgCl electrode (Kendall; Chicopee, MA, USA, 2.2 cm by 2.2 cm) was placed in the popliteal fossa, with the return electrode (Axelgaard; Fallbrook, CA, USA, 5 cm by 10 cm) placed just below the patella. Stimulation intensity was set to evoke a test (unconditioned) H-reflex below half maximum on the ascending phase of H-reflex recruitment curve to reduce the potential for evoking polysynaptic reflexes or self-facilitation (Hari *et al*., 2022). All H-reflexes were recorded at rest.

*H-reflex/M-wave recruitment curves* were collected from each participant by gradually increasing the TN stimulation, starting at an intensity that did not elicit an H-reflex or M-wave and increasing the TN stimulation until the maximum M-wave was achieved. H-reflexes were evoked every 5 seconds to minimize post-activation depression (Hultborn *et al*., 1996). The maximum peak-peak amplitude of the H-reflex and M-wave was used to calculate an Hmax/Mmax ratio for each participant. The average peak-to-peak amplitude of the test (unconditioned) H-reflex was expressed as a percentage of the maximum H-reflex ([test H / Hmax]*100%).

### Short-duration H-reflex suppression by antagonist CPN conditioning

In 23 participants with SCI (see Supplemental Table 1), the soleus H-reflex was conditioned by stimulating the ipsilateral common peroneal nerve (CPN) supplying the antagonist TA muscle (a.k.a. the common fibular nerve) as done previously in uninjured controls (Metz *et al*., 2023a). The CPN was stimulated using a bipolar arrangement (Ag-AgCl electrodes, Kendall; Chicopee, MA, USA, 2.2 cm by 2.2 cm) with the anode placed anterior and slightly distal to the fibular head on the leg. The cathode was placed near the fibular head in a location that elicited pure dorsiflexion. Three pulses (200 Hz, 1-ms pulse width) were applied to the CPN at an intensity of 1.0 and 1.5 x motor threshold (MT). Motor threshold was determined by the lowest CPN stimulation intensity that produced a discernable and reproducible M-wave in the TA muscle at rest of at least 30μV. Following a run of 7 unconditioned (test) soleus H-reflexes evoked every 5 s, 7 conditioned H-reflexes at one of the randomly chosen ISIs (3, 15, 30, 60, 100 or 200 ms) were elicited with 3-4 unconditioned test H-reflexes interposed between each run of conditioned H-reflexes to reestablish a steady baseline before the next set of conditioning stimuli.

#### Data analysis

The average peak-to-peak amplitude of the conditioned soleus H-reflexes at each ISI was compared to the average peak-to-peak amplitude of all the unconditioned (test) soleus H-reflexes evoked in a trial run. The % change H-reflex was expressed as: ([conditioned H-reflex – test H-reflex]/test H-reflex*100%). The % change at each ISI was then averaged across participants in each group. The effect of an isolated 1.0 and 1.5 x MT conditioning CPN stimulation on the soleus motoneurons was also measured in the resting soleus EMG. The rectified EMG from 7 to 10 CPN stimulation trials were averaged into 20 ms bins and the mean pre-stimulus EMG (or noise) measured between -100 ms and 0 ms was subtracted from each of the mean bin values. Bins containing EMG with CPN stimulation artifact (typically from 0 to 20 ms) were removed. The mean EMG in each time bin was then averaged across participants in each group. Data from 13 uninjured controls was used from the (Metz *et al*., 2023a) study.

### Long-duration H-reflex suppression by CPN and TN conditioning

To examine a longer time course of H-reflex suppression, soleus H-reflexes were conditioned with 1.5 x MT CPN stimulation as described above but at longer ISIs (500, 1000, 1500, 2000 and 2500 ms) in the same 23 participants with SCI (see Table 1) and compared to data from 13 controls from (Metz *et al*., 2023a). To examine if the long-duration suppression of soleus H-reflexes from CPN stimulation resembled the profile of repeated TN stimulation during RDD, in 34 participants with SCI (21 of whom also participated in the CPN experiment, Table 1) and in 16 controls (13 from Metz *et al*., 2022), repeated activation of the TN was examined at similar ISIs (500 to 2500 ms). During RDD, the first H-reflex (H1) of a stimulation trial of at least 10 H-reflexes was evoked just below 50% of maximum on the ascending part of the H-reflex recruitment curve (39.4±23.2% of Hmax). Each RDD interval was repeated 3 times.

#### Data analysis

The % change in the soleus H-reflex from the CPN stimulation applied at the longer ISIs was measured as above (% chg H-reflex_(CPN-TN)_). To quantify the amount of H-reflex suppression during RDD, the average peak-to-peak amplitude of the second to eighth H-reflex (H2-8) was expressed as a percentage of the averaged H-reflex (H1) using the formula: % chg H-reflex_(TN-TN)_ = [([avg H2-8] – H1) / H1]*100% for each stimulation frequency. In each participant, the resulting % chg H-reflex_(TN-TN)_ was averaged across the 3 trials for each RDD frequency and then averaged across participants in each group.

### Statistical Analysis

Statistical analysis was performed with Sigma Plot 11 software. The Shapiro-Wilk test was used to assess the distribution of the data with Student’s t-tests used to compare normally distributed data and Mann-Whitney Rank Sum test used to compare non-normally distributed data. The % change of the Ia-EPSPs or H-reflexes across the various ISIs during RDD was compared to a 0 % change using a one-way ANOVA or one-way ANOVA on Ranks, with various post hoc tests (e.g., Tukey, Holm-Sidak, Dunn’s Method) to determine which ISIs were significantly different from a 0% change (tests listed in Supplemental Tables). A two-way ANOVA was used to compare the % change H-reflex, Ia-EPSP or binned EMG values between groups (SCI, controls or drug/no drug) and ISI (or bin time) as factors. If there was a group and ISI interaction, the Holm-Sidak test was used to determine which ISI had a group difference. If there were no interaction effect, an unpaired Mann-Whitney Rank Sum or Student’s t-test was used with Bonferroni correction (see tests used in Supplemental tables). Data are presented in figures and in the text as mean and standard deviation (SD) except when standard error (SE) was used for easier visualization of the estimated spread of the binned EMG data. Significance was set as P <0.05 except for Bonferroni correction.

## Results

### Separate population of GABAergic neurons that innervate terminals and nodes of Ia afferents

#### Light-evoked GAD2 neurons

To explore whether GABA_B_ receptors on Ia afferents mediate presynaptic inhibition of the monosynaptic EPSP pathway to motoneurons, we took advantage of two distinct groups of GAD2+ GABAergic neurons with separate projections (Jankowska *et al*., 1981; Hari *et al*., 2022). One population projects to the dorsal nodes of Ia afferents to produce primary afferent depolarization (PAD) (Hari *et al*., 2022). The other population projects to the ventral horn and innervates the Ia afferent terminal located there (termed ventrally projecting GAD2 neurons), while at the same time innervating motoneurons in a triadic arrangement (Fig. 1A). The latter produces an IPSP on the motoneurons, which is a useful proxy for determining when these ventral GAD2 neuron terminals are activated. By employing GAD2//ChR2 mice with ChR2 in GABAergic neurons, we were able to activate this ventral projecting population fairly selectively by focusing laser light only on the motor nucleus (Fig. 1A-F), as seen by a ∼ 60 ms duration GABA_A_ receptor-mediated IPSP recorded on the motoneurons via ventral root (VR) recordings (Fig. 1C, Fig. 1E), and a relative lack of PAD measured from the dorsal roots (DR in Fig. 1B, Fig. 1F), indicating a lack of activation of the dorsal GAD2 neurons (Hari *et al*., 2022). In contrast, moving the laser dorsally (Fig. 1A-B) activated the dorsally projecting population of GAD2 neurons that produced a large PAD (Fig. 1B,F), but lacked any evoked GABAergic IPSP on motoneurons [Fig. 1E, as in (Hari *et al*., 2022)], again indicating that these dorsal GAD2+ GABAergic neurons innervate nodes of afferents and do not project to the motor nucleus.

Selective light activation of the ventral GAD2 neurons markedly inhibited the monosynaptic EPSPs evoked by Ia afferent stimulation 100 - 500 ms later (Figs. 1C-D), well after the IPSP (middle Fig. 1C) and action of any small amount of PAD that occurred (top Fig. 1C). This inhibition was absent in GAD2-mice and almost entirely blocked by the GABA_B_ receptor antagonist CGP55845 (bottom Fig. 1C-D). These findings suggest that the ventrally projecting GAD2 neurons produce presynaptic inhibition of the Ia terminal via GABA_B_ receptors that we have previously shown to be mainly located at the Ia terminal (Hari *et al*., 2022), potentially by inhibiting terminal VCa^+2^ channels and reducing neurotransmitter release.

Consistent with this conclusion, when recording directly from dorsal root afferents, the GABA_B_ receptor agonist baclofen caused a hyperpolarization of the afferent, as we detail later in the section “*Reduced GABA_B_ receptor action on post-activation depression and Ia hyperpolarization after SCI*” below. This hyperpolarization is possibly mediated by GIRK channels that are also activated by GABA_B_ receptors (Ciruela *et al*., 2010).

In contrast, light activation of the dorsal GAD2 neurons usually facilitated the EPSPs for < 100 ms, during the time period of phasic PAD (Fig. 1G,H), consistent with these neurons producing nodal GABA_A_ receptor-mediated facilitation of afferent transmission as previously detailed (Hari *et al*., 2022). To produce this facilitation in isolation, we took care to ensure the light intensity was not high enough to spread to the ventral GAD2 neurons to evoke an IPSP (Fig. 1E) or cause a PAD that was large enough to evoke spikes in the Ia afferents. Interestingly, GABA_B_ blockade enhanced the dorsal GAD2 neuron facilitation of the EPSP during the time window of PAD since, in theory, more neurotransmitter would be available to evoke the EPSP (Fig. 1H), without any stray activation of afferent terminal GABA_B_ receptors. Both PAD (Fig. 1F) and its facilitation of the EPSP (Fig. 1H) from dorsal light were blocked by the GABA_A_ receptor antagonist gabazine. When PAD-evoked spikes did occur, they often caused a monosynaptic activation of the test motoneurons, which caused post-activation depression of a subsequent test Ia-EPSP (Fig. 1I,J), an effect that lasted for several seconds as we previously reported (Metz *et al*., 2023a). Thus, dorsal GAD2 neurons can produce a PAD large enough to cause spiking of the Ia afferents that pre-activates the monosynaptic EPSP pathway prior to testing it, ultimately causing a GABA_A_-mediated post-activation depression of the EPSP that is distinct from presynaptic inhibition, though historically has been mistaken for presynaptic inhibition. This effect unexpectedly disappeared after SCI (Fig. 1J), detailed later.

Middle: DR-evoked EPSP without (pink) and with (blue) prior ventral light pulse, Bottom: DR-evoked EPSP without (blue) and with (green) CGP55845. **D)** Change in EPSP from prior ventral light pulse without and with CGP55845 (CGP) and in GAD2-mice. **E-F)** Amplitude of IPSP (E) and PAD (H) from ventral or dorsal light with and without gabazine. **G)** Dorsal (top) and ventral (bottom) root responses with (blue) and without (pink) PAD evoked from light pulse to dorsal spinal cord with DR-evoked EPSPs triggered during (left) and after (right) PAD. **H)** Change in EPSP when evoked during and after dorsal light PAD, and during PAD with and without CGP55845 and gabazine (Gbz) (447 nm laser, 1.1xLT). **I)** Same as in G but with stronger dorsal light pulse to produce PAD-evoked spikes in afferents (green) compared to no-spiking PAD (light blue, 1.2xLT). **J)** Difference in change in EPSP with PAD-evoked spikes from dorsal light pulse in control and SCI mice and in response to ventral light pulse. In D-J, * indicates p < 0.05 for paired and unpaired comparisons to the control variable; in H-J, + indicates p < 0.001 when the variable is compared to a 0% change or difference (see Supplemental Table 2A for mean (SD) values in D to J and results of statistical comparisons used).

#### Combined sensory and light activation of GAD2 neurons

Considering that GAD2 neurons are activated by spinal and supraspinal circuits, including the simple trisynaptic circuit from afferents to other afferents that underlies sensory-evoked PAD (Fig. 1A), we next examined the extent to which natural, sensory activation of GAD2 neurons modulate the Ia-EPSP by presynaptic inhibition produced from the ventral, GABA_B_ receptor-mediated pathway. Repeated activation of Ia afferents every 60 ms from a weak dorsal root stimulation (1.1 x T, Fig. 2A) caused a progressive increase in post-activation depression of the monosynaptic EPSP that likely engages the ventral, recurrent GAD2 neuron circuit (Fig. 2B pink). Optogenetic inhibition of the GAD2 neurons in GAD2//Arch3 mice (Fig. 2B green, C), or inhibition of the GABA_B_ receptor with CGP55845 (Fig. 2C), reduced this depression (RDD). These findings suggest that GAD2 neurons and associated GABA_B_ receptors cause presynaptic inhibition during post-activation depression produced during repeated Ia afferent activation. However, suppression of the Ia-EPSP from repeated stimulation was not completely eliminated by CGP55845, suggesting that the presynaptic mechanisms of terminal vesicle and neurotransmitter depletion (Sakaba & Neher, 2001a) continued to contribute to post-activation depression. Furthermore, repeated light activation of the ventral GAD2 neurons in GAD2//ChR2 mice when applied alone (Fig. 2F top blue) produced a similar amount of EPSP suppression as repetitive dorsal root activation when applied alone (Fig. 2D pink and Fig 2G), and also augmented suppression from repetitive dorsal root stimulation when applied with it (Fig. 2D blue and Fig 2G), the latter indicating that repetitive activation of sensory inputs alone does not engage the entire ventral GAD2 neuron pool. Although not completely eliminated, suppression of Ia-EPSPs was greatly reduced during GABA_B_ receptor blockade by CGP55845 during repetitive ventral light (Fig. 2F bottom blue) and dorsal root stimulation (Fig. 2E, top green), demonstrating a large GABA_B_ contribution to presynaptic inhibition during post-activation depression (arrows in Fig. 2G). The further suppression of Ia-EPSPs during the combined ventral light and dorsal root stimulation and its incomplete reduction by CPG55845 also demonstrates the amount of unused GABA_B_ effect on sensory evoked post-activation depression (arrow in Fig. 2G, right).

**Figure 2.**
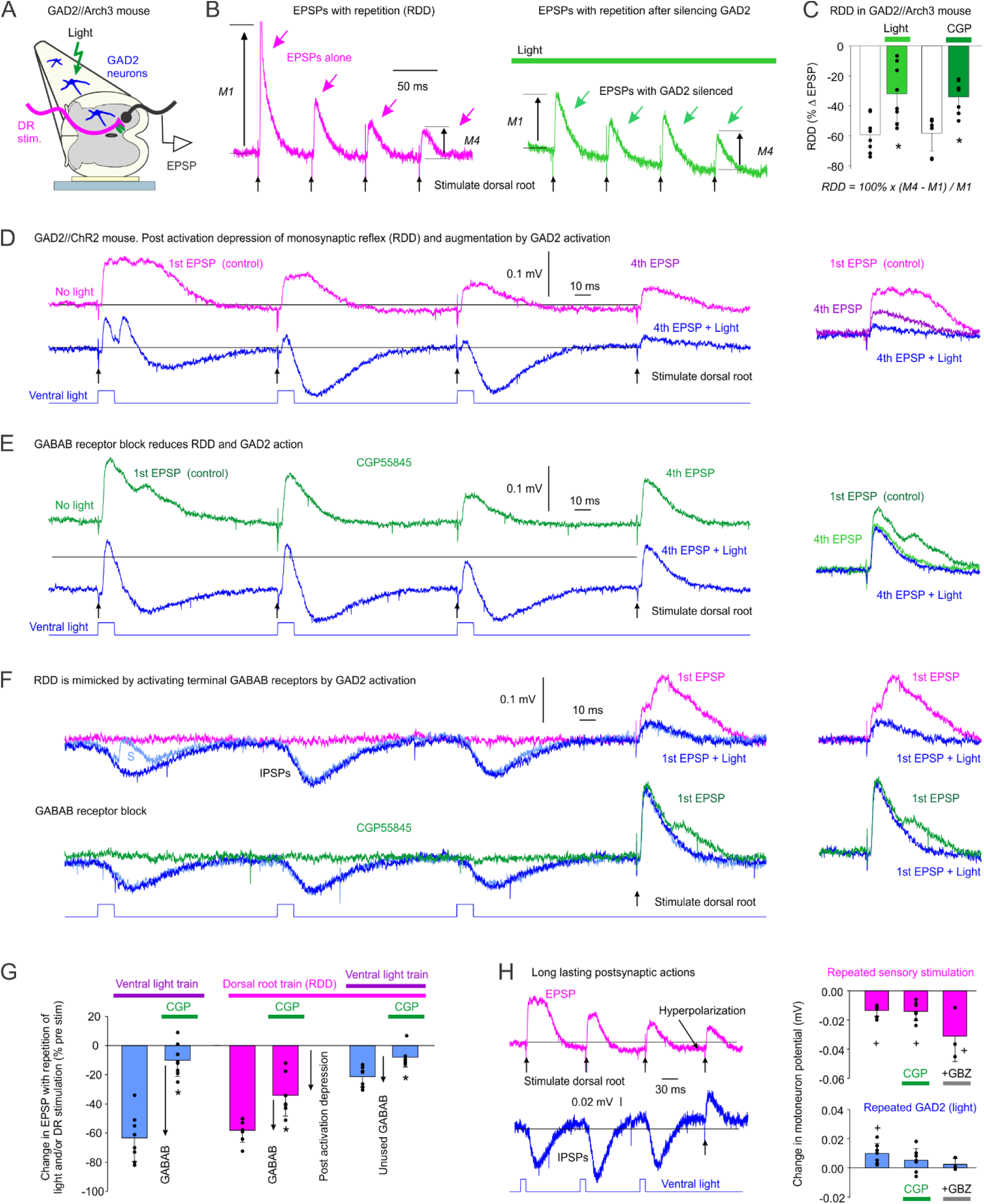
Post-activation depression, GAD2 neuron silencing and activation and GABA_B_ receptor blockade in mice. **A)** Schematic of dorsal root (DR) stimulation and ventral root (VR) recordings to produce post-activation depression of Ia-EPSPs. **B)** Evoked EPSPs from repetitive DR stimulation every 60ms in GAD2//Arch3 mice without (pink trace) and with (green trace) application of light to silence GAD2 neurons (532nm laser, 5 mW/mm^2^). Amplitude of 4^th^ EPSP (M4) is compared to first EPSP (M1). **C)** Percent change of 4^th^ EPSP compared to 1^st^ EPSP in GAD2//Arch3 mice (n = 8 VR recordings from 3 mice) without (white) and with (green) silencing of GAD2 neurons with light. Percentage change of EPSPs in wild type mice (n = 8 VR recordings in 3 mice) without (white) and with (green) GABA_B_ receptor blockade with CGP55845. **D-E)** Repetitive stimulation of Ia-EPSP from dorsal root stimulation alone (pink traces) and when combined with light pulses to ventral spinal cord (blue traces; 447nm laser, 0.7 mW/mm^2^, 1.5xLT)) in GAD2//ChR2 mice without (D) and during (E) GABA_B_ receptor blockade using CGP55845. **F)** Repetitive light stimulation of ventral root without (top) and with (bottom) CGP55845 before evoking a test EPSP with DR stimulation. Light blue trace (top) shows small EPSP from PAD-evoked spike. **G)** Change in Ia-EPSP size in response to repetitive dorsal root, ventral light or combined stimulation with and without CGP845 to indicate GABA_B_ receptor effect on post-activation depression. **H)** Baseline membrane potential of motoneuron in response to repetitive dorsal root stimulation (pink) or ventral light stimulation in GAD2//ChR2 mice without, and in presence of, CGP55845 and gabazine (GBZ). In C and G, * indicates p < 0.05 for paired Student t-tests and in H, + indicates p < 0.05 from a 0 mV change. Supplemental Table 2B contains mean (SD) values in C, G and H and results of statistical comparisons used.

While light activation of the ventral GAD2 neuron only caused a very small PAD, and rarely a delayed PAD-evoked spike, this did sometimes produce EPSPs on the motoneuron when the light intensity was increased between 1.1-1.5xT (Fig. 2F light blue trace, with EPSP labelled s), suggesting that the ventral GAD2 neurons may innervate GABA_A_ receptors on some afferents, but not necessarily the Ia afferents. However, this did not enhance post-activation depression of the monosynaptic EPSPs (Fig. 1J), indicating that the axons activated by the ventral GAD2 neuron in this way do not include the Ia afferents in the tested monosynaptic EPSP pathway (Fig. 1A), and instead could include higher threshold Ia afferents, other rapid group Ib or II afferents or even interneuron axons.

Thus, overall post-activation depression from repetitive Ia afferent activation is a complex, three-part process that involves ventral and dorsal GAD2 neurons that activate GABA_B_ and GABA_A_ receptors on afferents respectively, with the latter causing both nodal facilitation and PAD-evoked spikes. Finally, we noticed that even at long intervals up to 500 ms, repeated dorsal root stimulation caused a gradually increasing baseline hyperpolarization of motoneurons (Fig. 2H pink) that persisted during GABA_B_ or GABA_A_ receptor blockade. This small baseline hyperpolarization is unlikely to change the Ia-EPSP but should inhibit motoneuron spike initiation and associated H-reflexes in the awake state, adding a fourth mechanism underling the increase in post-activation depression from repetitive Ia activation. In contrast, repeated ventral light stimulation produced a small increase in baseline depolarization (Fig. 2H blue) that was reduced with both GABA_A_ or GABA_B_ receptor blockade. Combined these results indicate that spinal circuits activated by natural sensory activation can also produce long-lasting postsynaptic inhibition of the motoneuron that may contribute to post-activation depression, at least up to 500 ms.

### Reduced GABA_B_ receptors on Ia afferent terminals after SCI

Because presynaptic inhibition of the Ia afferent is affected by GABA_B_ receptor activation and is reduced after SCI, we wondered whether this reduction is associated with a reduction of GABA_B_ receptors on the Ia afferent terminal (Fig. 3A). To examine this, sensory afferents were labelled in the dorsal root and spinal cord of adult GAD2//ChR2-EYFP mice by a peripheral adeno-associated virus (AAV9-tdTom) injection, as shown from a confocal image of transverse spinal cord sections (Fig. 3E intact, Fig. 3F SCI, top panels). The large myelinated proprioceptive Ia afferents were identified by characteristic extensive ventral horn branching and unique innervation of motoneurons and with their terminals being VGLUT1+ (pink Fig. 3E,F, 4th row). Overall, we found that with SCI the expression of GABA_B_ receptors on these Ia afferent terminals were reduced by about half (Fig. 3E vs 3F, 5^th^ row), with a lower receptor intensity in terminals (Fig. 3B left). In contrast, the GABA_B_ receptor intensity in motoneurons did not change (Fig. 3B right), and thus, the GABA_B_ receptor intensity in the Ia terminals relative to that in the motoneurons was also reduced with SCI (Fig. 3C). The number of Ia terminals with receptors did not change with SCI (Fig. 3D).

**Figure 3.**
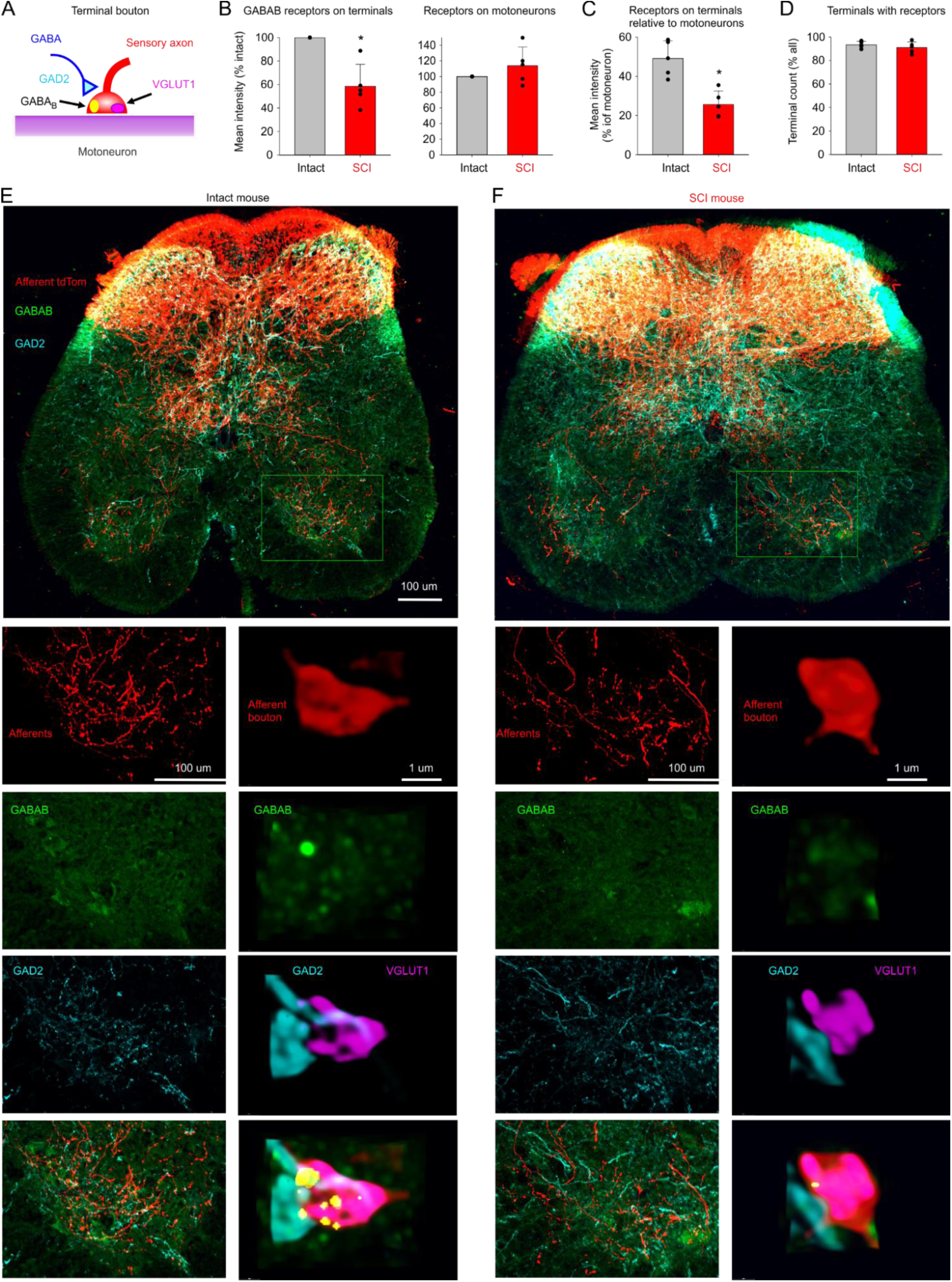
Reduced GABA_B_ receptors on Ia afferent terminals after SCI. **A)** Schematic of immunolabelling of Ia afferents and GAD2+ interneuron terminal and associated GABA_B_ receptors and VGLUT1+ afferent terminals. **B)** Left: Mean intensity of GABA_B_ receptors on the Ia terminal of SCI mice as a % of the intensity in intact mice, with pairs of injured and control spinal cord sections placed on the same slide for labelling. Right: Mean intensity of GABA_B_ receptors on the motoneuron of SCI mice as a % of the intensity on the motoneurons of control mice. **C)** Mean intensity of GABA_B_ receptors on the Ia terminal as a % of the intensity on the motoneuron for SCI and control mice. **D)** Mean number of terminals with GABA_B_ receptors as a % of all identified Ia terminals. **E-F)** Transverse section of spinal cords from intact (**E**) and chronic SCI (**F**) mice whose afferents were labeled with tdTOM (red; AAV9-tdTom), GABA_B_ receptors in green and GAD2 interneurons in blue in GAD2//ChR2-EYFP mice (top row). Lower left panels showing enlarged images of afferents (2nd row), GABA_B_ receptors (3rd row), GAD2 neurons (4th row) and merged picture from all three (5^th^ row). Right panels: expanded view of Ia afferent terminal for data on left with additional labelling of Ia terminal with VGLUT1 in 4th and 5^th^ row. GABA_B_ receptors computed to be in the volume of the terminal are rendered yellow. N = 5 mice per group. * p < 0.05 for unpaired Student’s t-tests. Mean (SD) and statistical results in Supplemental Table 2C.

### Reduced GABA_B_ receptor action on post-activation depression and Ia hyperpolarization after SCI

Since the GABA_B_ receptor expression on the Ia terminal is reduced after chronic SCI and GABA_B_ partly mediates post-activation depression (RDD), the previously reported reduction of post-activation depression with SCI may be due to a reduction of GABA_B_ receptor action. Thus, we next examined this by blocking GABA_B_ receptors with CGP55845 during post-activation depression. We first characterized the rate-dependency of post-activation depression (i.e., RDD) of the Ia-EPSP in control mice at ISIs from 0.5 to 5 s to match the rates we used for H-reflexes in human participants presented in the section “*Reduced post-activation depression with repeated reflex testing in humans with SCI*” below. In control mice, Ia-EPSPs were reduced at all ISIs (< 0% change, Fig. 4A black), with a suppression > 40% when dorsal root stimulation was applied repetitively every 0.5 and 1 s. Overall, Ia-EPSP suppression was reduced to < 40% in mice with chronic SCI (Fig. 4A pink), and was less compared to control mice, specifically at the 0.5 and 1 s ISIs. Application of CGP55845 to the spinal cords of intact control mice reduced the suppression of the Ia-EPSP to levels measured in SCI mice (Fig. 4B blue), specifically at the 0.5 and 1 s ISIs. In contrast, CGP55845 did not reduce post-activation depression in mice with SCI (Fig. 4C, blue), with instead a small increase at the 0.5 ms ISI. This lack of a consistent decrease of RDD following CGP55845 after SCI supports the conclusion that decreased GABA_B_ receptors leads to less presynaptic inhibition and RDD after SCI.

**Figure 4.**
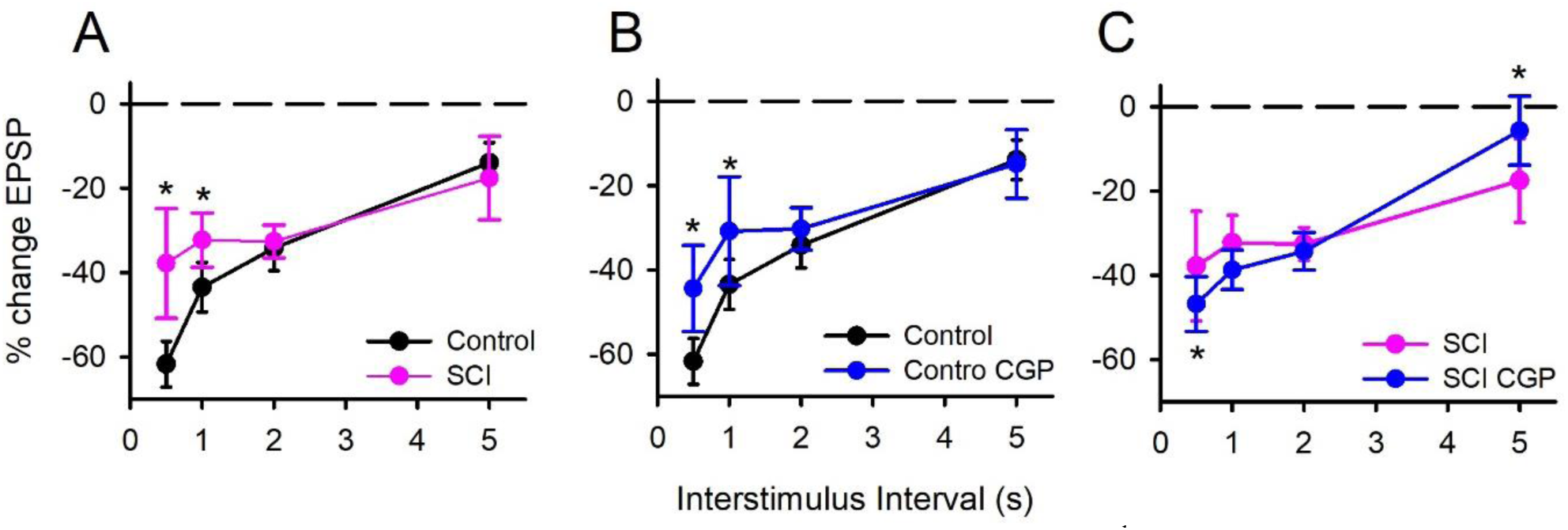
Rate-dependent depression of Ia-EPSPs. **A)** % change of 4^th^ Ia-EPSP as a % of 1^st^ Ia-EPSP at 0.5, 1, 2 and 5 s ISIs in control (black) and SCI (pink) mice (n = 3 mice per group). **B&C)** % change in Ia-EPSP in control **(B)** and SCI **(C)** mice before and after bath application of CGP55845 (blue). Results of statistical tests in Supplemental Table 3. * p ≤ 0.005. We next examined in intact control mice if facilitation of GABA_B_ receptors on afferents by the agonist baclofen has less action in SCI compared to intact mice, consistent with their lower GABA_B_ receptor expression in Ia afferent terminals. To do this we recorded the DC membrane potential of the cut central portion of a dorsal root to get a compound afferent potential from large afferents (Fig. 5 left) (Hari *et al.,* 2022). Recordings were made during blockade of fast synaptic transmission to synaptically isolate the axons before adding baclofen (50 μM CNQX, APV and gabazine, and 5 μM strychnine). In intact control mice (Fig. 5 top left trace), baclofen produced a hyperpolarization of the sensory afferents, consistent with GABA_B_ receptors facilitating downstream, outward GIRK channels (Luscher & Slesinger, 2010). The amount of hyperpolarization from baclofen was reduced in chronic SCI mice (Fig. 5 bottom left trace), compared to the control (normal) mice (Fig. 5 right).

**Figure 5.**
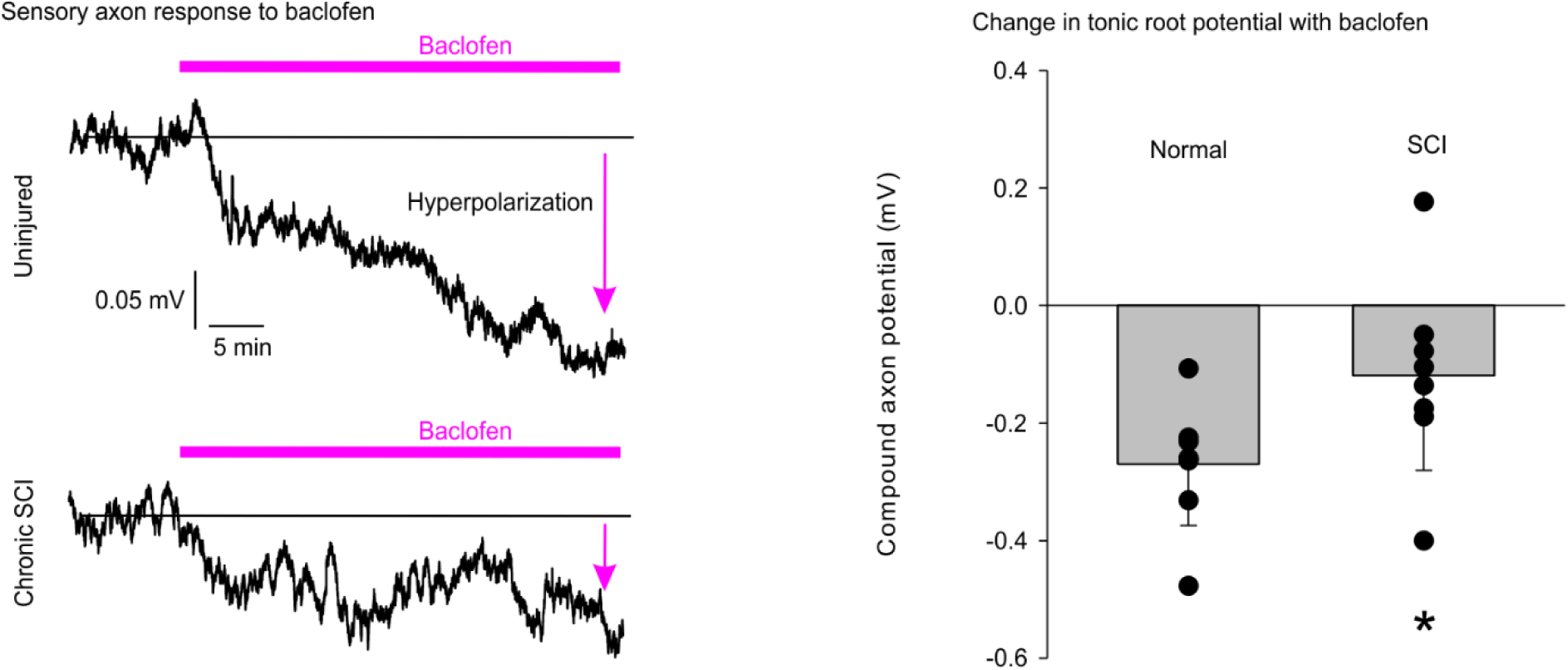
Hyperpolarization of Ia afferents by baclofen. Left: Sacral (S4) dorsal root potential recording from a normal (top) and chronic SCI (bottom) mouse starting from a stable baseline in presence of synaptic blockade. Horizontal pink line indicates addition of baclofen to the bath. The amplitude of the baclofen-induced hyperpolarization from baseline is marked by pink arrows. **Right:** Mean (SD) difference between the steady dorsal root potential (in mV) before and during baclofen application in normal and SCI mice. Overlaid individual data points (circles), data from 6 dorsal roots recorded from two normal mice and data from 6 dorsal roots recorded from two mice with chronic SCI. * indicates p = 0.044, unpaired Student’s t-test (see Supplemental Table 2D for mean (SD) values and full results of statistical tests.

In summary, the reduced effect of the GABA_B_ receptor antagonist CGP55845 on post-activation depression and the reduced hyperpolarization of Ia afferents by facilitating the GABA_B_ receptors with baclofen after SCI are both consistent with a reduced number of GABA_B_ receptors on the Ia afferent terminals as demonstrated in Figure 3 above.

### Reduced post-activation depression with repeated reflex testing in humans with SCI

Using post-activation depression as a possible proxy for changes in GABA_B_ receptor action after SCI, we next examined whether post-activation depression evoked by repeated reflex testing was reduced in humans with spinal cord injury, like in mice. Post-activation depression of the mainly monosynaptic H-reflex in humans was examined in the soleus muscle in response to repeated tibial nerve (TN) stimulation, with control H-reflexes just below half maximum to allow optimal sensitivity to changes (red arrows in Fig. 6A). Repetitive stimulation of the TN at increasing rates (ISIs between 5 to 0.5s) produced increasing depression of the H-reflex in control participants, consistent with a marked post-activation depression that was dependent on stimulation rate (rate dependent depression, RDD). This RDD was reduced in participants with chronic SCI (pink, Fig. 6C) compared to controls, specifically at the 0.5, 1 and 2 s ISIs. While the absolute values of both the H_max_ and M_max_ were smaller in the SCI participants (Fig. 6Bi), the mean H_max_/M_max_ ratio was not different compared to controls, with ratios ∼50% in both groups Fig. 6Bii), suggesting that the reduction in reflex depression with SCI was not due marked changes in the proportion of the motoneuron pool recruited by the reflex.

**Figure 6:**
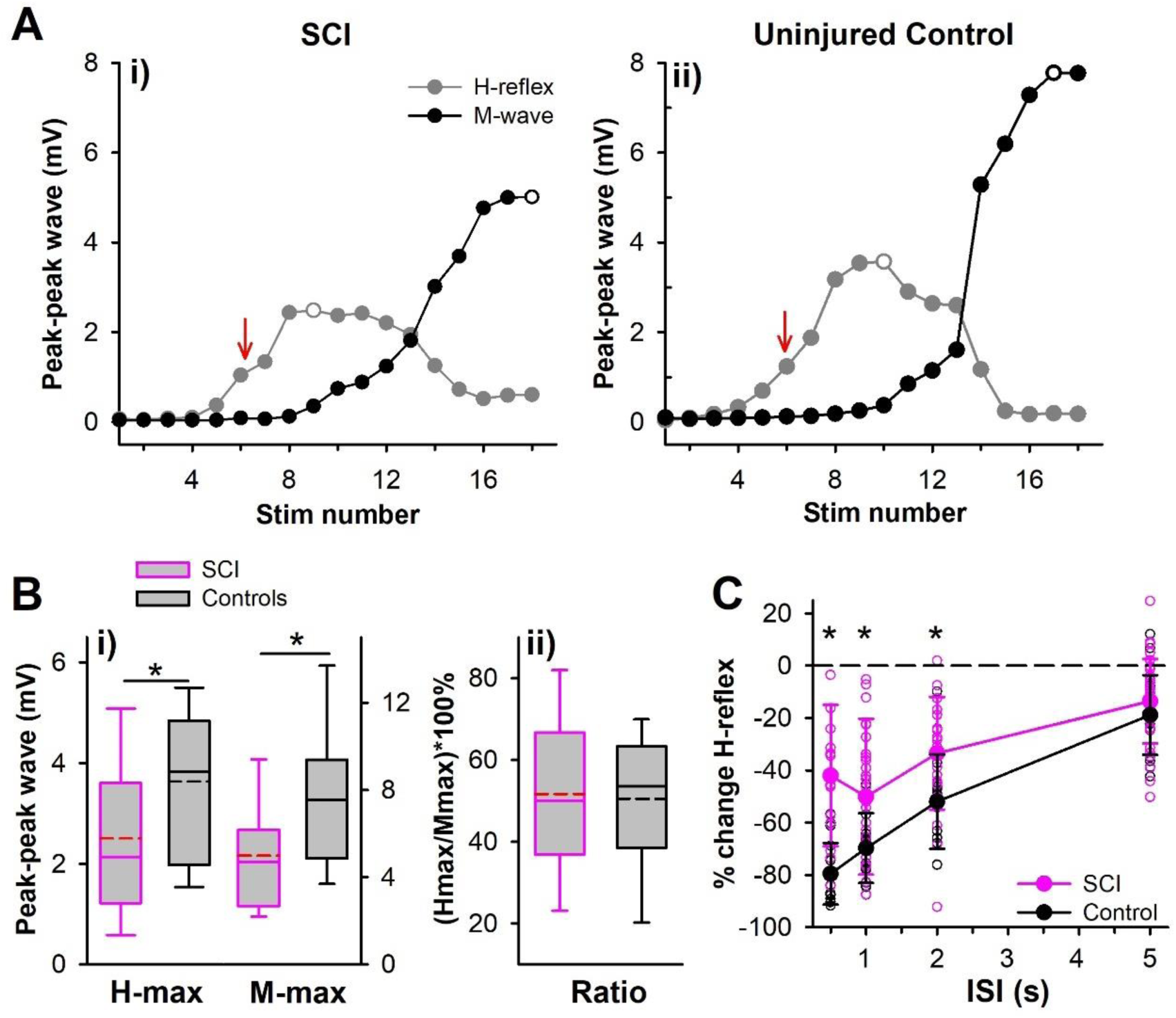
H-reflex/M-wave recruitment curves. **A)** Peak-to-peak amplitude of the soleus H-reflex (grey circles) and M-wave (black circles) from a participant **(i)** with SCI and **(ii)** a control plotted against stimulation number during increasing TN stimulation. Maximum H-reflex and M-wave marked with open circles and amplitude test H-reflex used in subsequent conditioning experiments marked by red arrows. **B)** Box plot of maximum H-reflex and M-wave **(i)** and H_max_/M_max_ ratio **(ii)** with mean (dashed) and median (solid) lines, 25th and 75th percentiles by the box bounds, and the 95th and 5th percentiles by whiskers. **C)** Mean % change of soleus H-reflex (large circles) from repetitive TN stimulation in the SCI (n=34) and control [n=16, data taken from (Metz *et al*., 2023a)] groups plotted as a function of the interstimulus interval (ISI), individual means plotted with small circles. Whiskers = SD, * = P < 0.05. Mean (SD) and results of statistical tests in Supplemental Table 2D for data in B and Supplemental Table 5 for C.

### Reduced suppression of soleus H-reflexes by antagonist CPN stimulation in SCI

Repeated reflex testing, as detailed above (Fig. 6), produces a post-activation depression (RDD) that is dominated by repeated activation of the synapse in the monosynaptic Ia-EPSP pathway and associated transmitter depletion, and to a lesser extent, activation of GABA_B_ receptors caused by indirect recruitment of GABAergic systems from the repeated afferent stimulation as shown in the mice experiments of Figure 2. Thus, to study the action of GABA with a potentially smaller amount of post-activation depression (and neurotransmitter depletion), we next turned to stimulating the antagonist nerve (CPN) prior to testing the soleus H-reflex, which is known to activate the GABAergic neurons in the soleus reflex pathway that should produce both presynaptic inhibition and PAD [Figs. 1-2, see also (Hari *et al*., 2022; Metz *et al*., 2023a)]. At short intervals this conditioning CPN action can still be complex, as detailed in mice for direct activation of GABAergic GAD2 neurons, involving both GABAergic activation of Ia afferent terminals to produce GABA_B_ receptor-mediated presynaptic inhibition and activation of nodes to produce GABA_A_ receptor-mediated PAD. The PAD in turn has two opposing actions, increasing Ia afferent conduction by nodal depolarization at short conditioning intervals during phasic PAD and decreasing transmitter released from terminals over longer intervals due to PAD-evoked spikes in the soleus Ia afferents that produce post-activation depression (depression of transmitter release and GABA_B_ action), though the latter action dominates when the conditioning stimulation is large, as we used here. Thus, we next examined how antagonist-evoked depression of the soleus H-reflex changed with spinal cord injury as an indicator of changes in GABA mediated inhibition, specifically focusing on larger conditioning intensities that promote inhibition. To again assure that a similar group of motoneurons were tested with and without SCI, the average amplitude of the test H-reflex used in the SCI participants was noted to be 27.5 (21.3)% of H_max_ and not different to that used in controls at 30.9 (13.8) % (p = 0.300, unpaired Student’s t-test). We first examined short ISIs between 3 and 200 ms at a low [1.0 x CPN motor threshold (MT), Fig. 7Ai, ii] and higher [1.5 x MT, Fig. 7Ci, ii] intensity of conditioning stimulation as done previously (Metz *et al*., 2023a). As detailed previously in control participants and mice, after SCI the CPN stimulation evoked a paradoxical early soleus response (non-classical reflex), with a short latency (∼ 35 ms), consistent with it being caused by PAD-evoked spikes in the soleus Ia afferents that then directly activated the test soleus motoneurons. The lower conditioning stimulation intensity of the CPN produced an early soleus response that was not smaller than in control subjects (Fig. 7Bii), with responses in both groups increasing with CPN intensity (from 1.0 to 1.5 x MT; Fig 7Aiii and Ciii), and with stronger responses showing less confounding facilitation of the H-reflex, as detailed above, suggesting that after SCI, PAD still readily evoked spikes in Ia afferents to the tested muscle. During weak conditioning CPN stimulation at 1.0 x MT, H-reflex suppression was greater in the control group compared to the SCI participants across all ISIs, especially clear at the 30 ms ISI. At the stronger 1.5 x MT conditioning CPN stimulation, the early soleus response was larger in the SCI participants compared to controls (Fig. 7Dii), potentially reflecting a larger number of PAD-evoked spikes. Despite this, the amount of H-reflex suppression in the SCI participants remained less compared to the controls (Fig. 7Di), specifically at the 100 ms ISI. At both conditioning CPN stimulation intensities, modulation of the soleus H-reflex did not follow the profile of the early soleus response (Fig. 7B and D), ruling out a large postsynaptic influence on the H-reflex. Overall, SCI decreased the CPN-evoked depression of the H-reflex, despite putative maintenance or even augmentation of GABA_A_ mediated PAD actions (spikes), pointing toward a decrease in GABA_B_ receptor-mediated presynaptic inhibition of the monosynaptic reflex with SCI, separate from post-activation depression caused from transmitter depletion.

**Figure 7:**
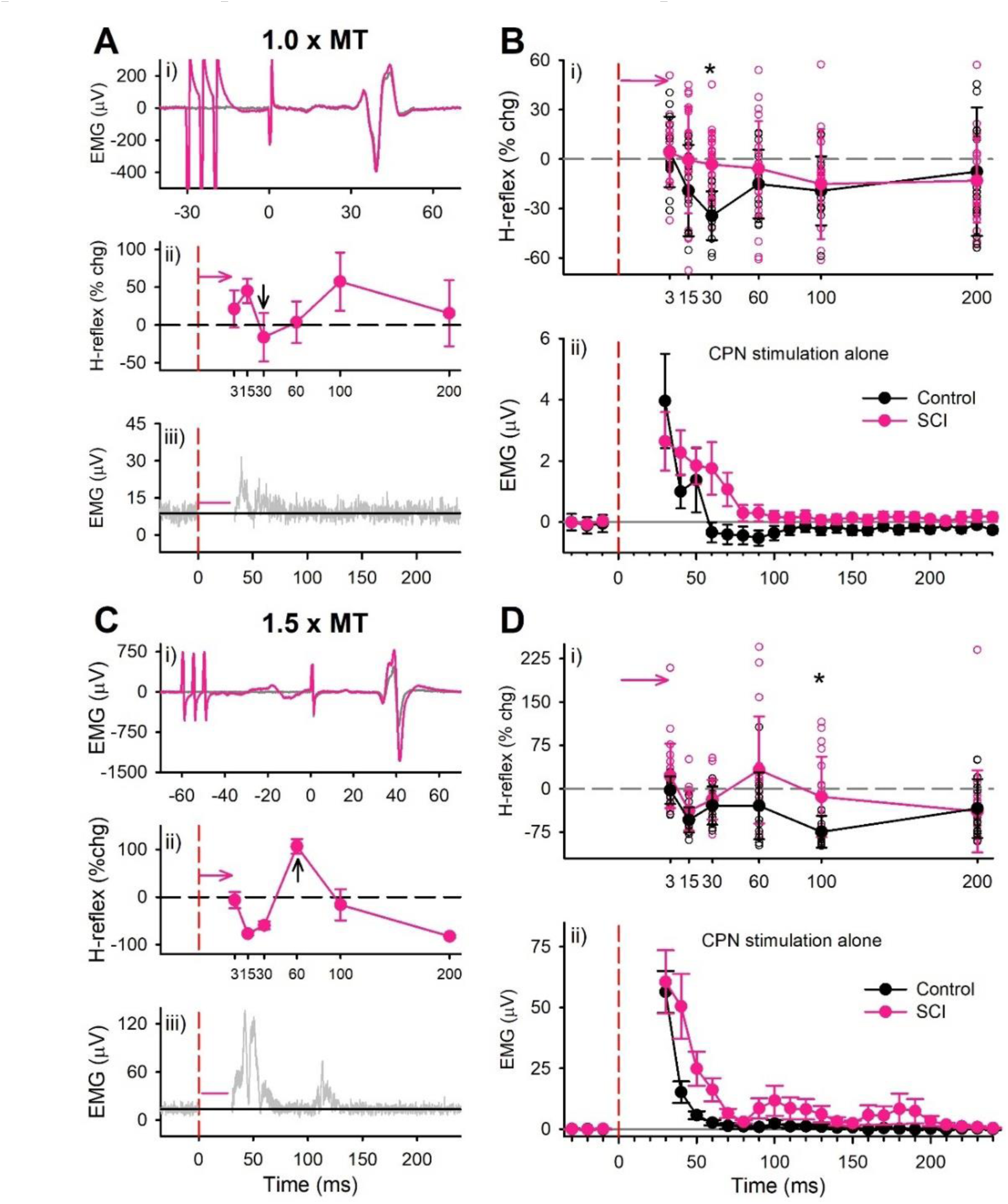
Conditioning soleus H-reflex by CPN stimulation at short ISIs. A,C) Representative data from a participant with SCI at the 1.0 x MT (**A**) and 1.5 x MT (**C**) stimulation intensities. **i**) Average of 7 conditioned (pink) and 7 test (grey) soleus H-reflexes (unrectified EMG) at the 30 ms (**A**) and 60 ms (**C**) ISI at rest. TN stimulation to evoke H-reflex applied at 0 ms. **ii)** Mean (±SD) % change of the H-reflex plotted at each ISI. Time of CPN stimulation marked by red dashed line. H-reflexes are shifted to the right of the time of CPN stimulation by the latency of the H-reflex (∼30 ms, (Metz *et al*., 2023b)). **iii)** Average rectified soleus EMG (8 sweeps) measured at rest with CPN stimulation delivered alone. Stimulation artifact and TA M-wave cross talk has been removed as marked by pink horizontal line. **B,D)** Group data. **i)** Mean (±SD) % change in soleus H-reflex averaged across the SCI (pink, n=23) and control (black, n=13) groups for the 1.0 x MT (**B**) and 1.5 x MT (**D**). CPN intensities. Small circles indicate, for individual participants, the average of the 7 conditioned H-reflexes at each ISI as a percentage of all test H-reflexes evoked in the trial. **ii)** Mean (±SE) soleus EMG from CPN stimulation alone at rest, values averaged into 10 ms bins with the mean, pre-stimulus value (100 ms window just prior to stimulation) subtracted from each bin. * = significant difference between the SCI and control group (p < 0.05). Results of statistical tests in Supplemental Table 4.

### The early soleus response and H-reflex suppression

Since the early soleus response following the CPN conditioning may reflect the activation of PAD-evoked spikes that depress the H-reflexes via post-activation depression, we expect that a larger number of afferents with PAD-evoked spikes may produce a larger suppression of subsequent soleus H-reflexes. In support of this, in controls there was a significant correlation when plotting the amplitude of the early soleus response to CPN stimulation (between 30 and 50 ms post CPN stimulation) and the amount of soleus H-reflex suppression at the 100 ms ISI (Fig. 8, right graph: r = -0.84). A similar significant correlation was found for the participants with SCI (Fig. 8, left, r = -0.57). However, the slope of the straight line fit through the data was 2.5 times as shallow in the SCI participants (-0.69 %chg H-reflex/μV early soleus response) compared to controls (-1.74 %chg H-reflex/μV early soleus response). This suggests that the effects on presynaptic inhibition of the soleus Ia afferents from the activation of GABA neurons by the conditioning CPN stimulation had a weaker effect in participants with SCI compared to controls, and again this might result from weaker GABA_B_ receptor-mediated presynaptic inhibition of the Ia afferent terminals (from conditioning-evoked GABA release on the Ia terminals), despite the PAD and PAD-evoked spikes not being smaller with SCI, as detailed in mice.

**Figure 8.**
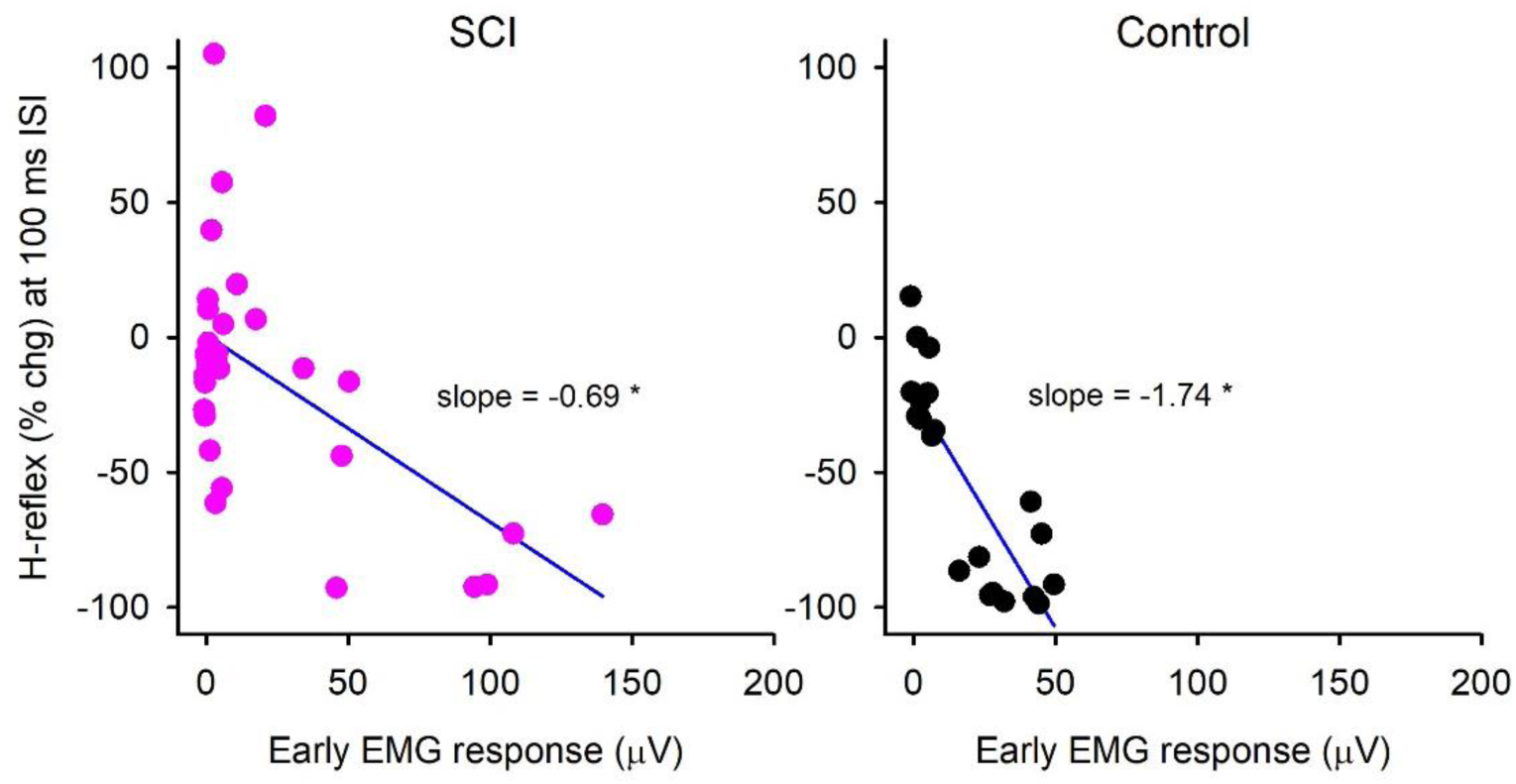
Relationship between conditioned H-reflex suppression and early reflex EMG. Percentage change (% chg) in the conditioned SOL H-reflex at the 100 ms ISI plotted against the resting soleus EMG 30-50 ms after the CPN stimulation (early EMG response) for participants with SCI (left) and controls (right). There was 34 data points from 20 participants in the SCI data (left) with 6 early EMG points unavailable and 22 data points from 13 participants in the control data (right) with 4 early EMG points unavailable. Data from the 1.0 and 1.5 x MT stimulation are combined. Pearson’s product moment correlation (r) was significant at p<0.001 (marked with *).

### Longer interval soleus H-reflex suppression from CPN and TN conditioning

A conditioning afferent (CPN) stimulation has less complex action at long intervals, where confounding GABA-evoked PAD no longer facilitates afferent conduction, and inhibition caused by presynaptic GABA actions dominate. Thus, we next examined if the suppression of the soleus H-reflex from antagonist CPN conditioning lasted for several seconds, as would be expected from a long-lasting presynaptic inhibition of the Ia afferents caused by GABA_B_ receptor action (Figs. 1 and 2) and transmitter depletion (Sakaba & Neher, 2001b), which causes a subsequent post-activation depression of the soleus Ia-EPSP (Hultborn *et al*., 1996). As we have previously demonstrated in controls (Metz *et al*., 2023a), the amplitude of the soleus H-reflexes was reduced by a 1.5 x MT CPN conditioning stimulation for many seconds (Fig. 9A, black squares), with the conditioned H-reflexes below a 0% change at all ISIs out to 2.5 s (open squares). The CPN-conditioned H-reflexes were also suppressed in the SCI participants (Fig. 9A, pink squares) but only out to the 2 s ISI (open circles). Overall, the suppression of the CPN-conditioned H-reflexes was greater in controls compared to the SCI group, specifically at the 0.5 and 1 s ISIs.

**Figure 9:**
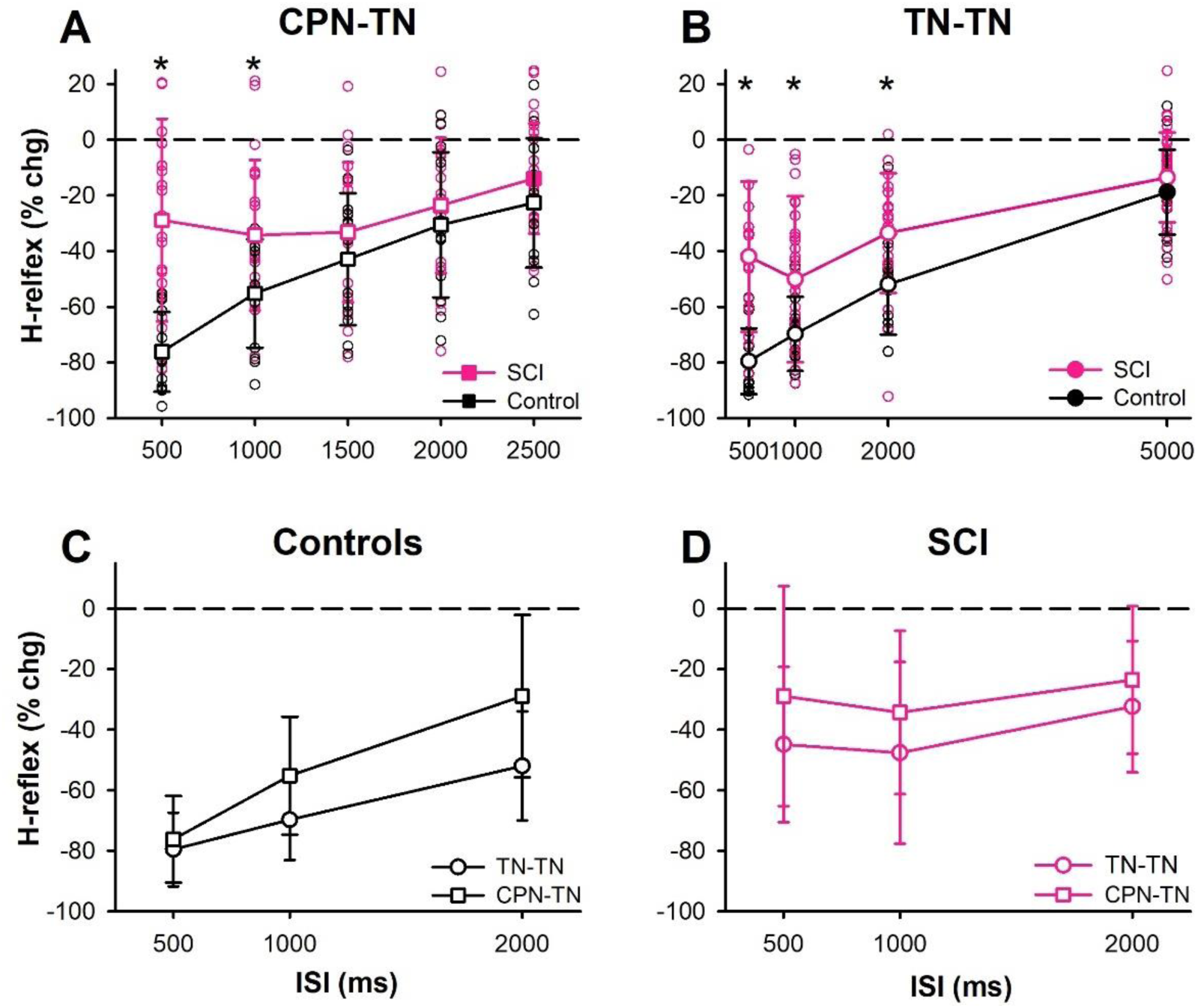
Slow RDD of soleus H-reflex. **A)** Mean (±SD) % change of soleus H-reflexes from a 1.5 x MT CPN conditioning stimulation (CPN-TN) in the SCI (pink, n=23) and control (black, n=13) groups plotted as a function of ISI. **B)** Mean (±SD) % change of soleus H-reflex from repetitive TN stimulation (TN-TN or RDD) in the SCI (n=34) and control (n=16) groups plotted as a function of ISI, modified from Fig. 5C. **C & D)** Same data as A and B but plotted to compare CPN conditioning and RDD within each group. White symbols mark conditioned H-reflexes that are significantly different than a 0% change. Small circles show individual data points. * indicates difference between groups at a given ISI. Results of one and two-way ANOVAs and post-hoc comparisons in Supplemental Table 5.

The amplitude and pattern of H-reflex suppression from CPN conditioning, and the reduction after chronic SCI, was similar to that produced when the soleus Ia afferents were directly activated by repeated TN stimulation (Fig. 9B, data replotted from Fig. 5C), suggesting that both forms of conditioning (CPN-TN and TN-TN) were mediated by post-activation depression. When comparing the amount of H-reflex suppression during repeated TN stimulation (TN-TN) to that during CPN conditioning (CPN-TN) within the control (Fig. 9C) or SCI (Fig. 9D) groups, as expected H-reflex suppression was larger during repeated TN stimulation compared to the CPN conditioning, but not at any specific ISI in either group.

### Baclofen increases post-activation depression in SCI

Finally, to more directly test whether GABA_B_ receptors are involved in presynaptic inhibition in humans, we examined the action of the GABA_B_ receptor agonist baclofen in SCI participants taking oral baclofen. Because GABA_B_ receptors inhibit Ca^2+^entry into the presynaptic terminal, baclofen should inhibit both transmitter release and vesicle replenishment, both of which are Ca^2+^-dependent (Sakaba & Neher, 2001a). Thus, baclofen should theoretically increase post-activation depression caused by transmitter depletion with repeated activation by reducing the readily releasable pool of vesicles [as shown in (Sakaba & Neher, 2003) for the Calyx of Held], whereas the GABA_B_ antagonist CGP55845 should decrease post-activation depression by blocking GABA_B_ receptors that inhibit both transmitter release and vesicle supply (Figs. 1 and 2) [although see (Orsnes *et al*., 2000)]. To examine baclofen action, we compared the magnitude of H-reflex suppression with repeated TN stimulation between SCI participants that were not taking baclofen with participants who were taking daily oral baclofen (> 30 mg/day, Supplemental Table 1). As expected, in participants not taking oral baclofen, when H-reflexes were repetitively activated every 1 s (Fig. 10A), the last 5 H-reflexes in the stimulus train were less suppressed compared to participants taking daily oral baclofen and to the uninjured control group (Fig. 10B). No differences in H-reflex suppression between participants taking and not taking baclofen were observed at the longer ISIs of 2 or 5 s (p > 0.436, unpaired Student’s t-test, data not shown), which is outside of the GABA_B_ receptor effect as shown in mice. Although baclofen also decreases the H-reflex directly via its inhibition of transmitter release, this was not tested, and the dose used in participants did not completely eliminate reflexes as can happen during intrathecal baclofen application (Stokic *et al*., 2006).

**Figure 10:**
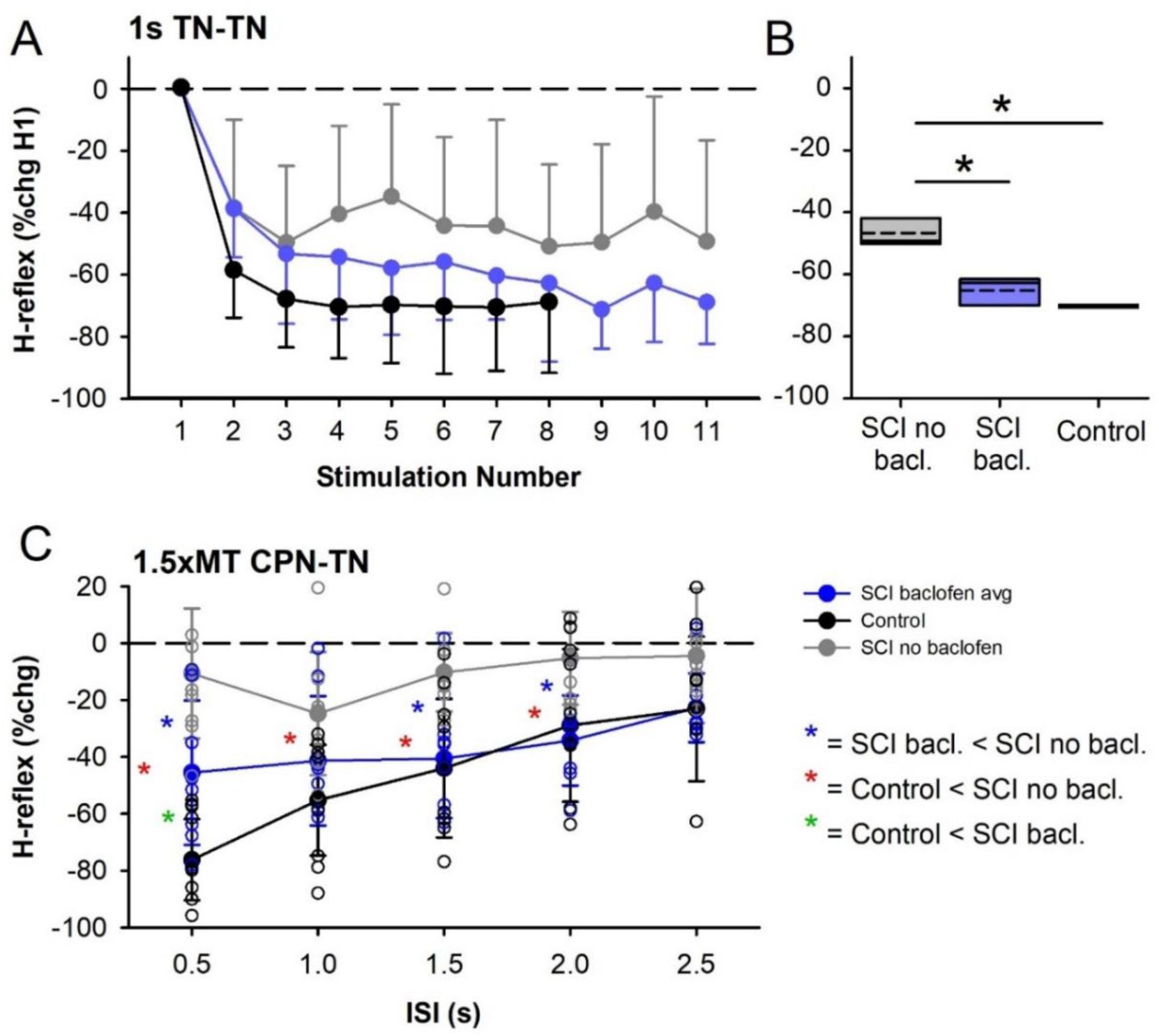
Daily oral baclofen increases post-activation depression in SCI. **A)** Mean (SD) % change of H-reflexes compared to the 1^st^ H-reflex during repetitive TN stimulation every 1 s in SCI participants not taking baclofen (grey, n = 8), SCI participants taking baclofen (blue, n = 6) and uninjured control participants [black, n = 7 from (Metz *et al*., 2023a)]. **B)** Box plot of average of 6^th^ to 11^th^ H-reflex in SCI participants and 4^th^ to 8^th^ H-reflex in control participants, with mean (dashed) and median (solid) lines and 25th and 75th percentiles by the box bounds. **C)** Mean (SD) % change of H-reflex conditioned by a 1.5 x MT CPN stimulation at various ISIs in SCI participants not taking baclofen (n = 9), SCI participants on baclofen (n = 8) and uninjured controls [n = 13 from (Metz *et al*., 2023a)]. Small symbols show individual participant values. Same SCI participants as in B plus a few additional with CPN-TN data. Mean (SD) values for **B** and results of all statistical tests in Supplemental Table 6.

Similar to H-reflex suppression from repetitive TN stimulation, the suppression of soleus H-reflexes from a prior conditioning CPN stimulation at ISIs from 0.5 to 2.5 s was less in the SCI participants not taking baclofen compared to those taking baclofen and to the uninjured controls (Fig. 10C). Specifically, H-reflex suppression in SCI participants not taking baclofen was less compared participants on baclofen at the 0.5, 1.5, and 2 s ISIs (Fig. 10C *) and likewise less suppressed compared to controls from the 0.5 to 2 s ISIs (Fig. 10C *), with the baclofen group having lower H-reflex suppression compared to controls only at the 0.5 s ISI (Fig. 10C *).

## Discussion

Our results provide evidence that a distinct population of ventrally projecting GAD2 neurons activate GABA_B_ receptors on Ia afferent terminals to mediate presynaptic inhibition of Ia afferents and this contributes to post-activation depression of Ia-EPSPs during repeated stimulation (RDD). Consistent with their triadic innervation of Ia afferent terminals, these ventral GAD2 neurons also produce a short GABA-mediated IPSP on motoneurons that is useful to confirm the release of GABA onto the terminal region and distinguish these GAD2 neurons from dorsal GAD2 neurons that do not produce this IPSP. This ventral GAD2-mediated presynaptic inhibition is largely blocked by the GABA_B_ receptor antagonist CGP55845, further disproving the classical idea that GABA_A_ receptors are involved in presynaptic inhibition, and consistent with a lack of GABA_A_ receptors or their action on these Ia afferent terminals (Hari *et al*., 2022). We propose that activation of homonymous Ia afferent collaterals activate first order neurons that specifically activate these ventrally projecting GAD2 neurons that in turn activate GABA_B_ receptors on mostly homonymous Ia afferent terminals to reduce Ca^2+^ flow into the presynaptic terminal, neurotransmitter release and vesicle replenishment during repetitive stimulation. In contrast, a separate population of more dorsally projecting GAD2 neurons seem to account for classic GABA_A_ receptor-mediated PAD in Ia afferents (and not motoneuron IPSPs), which functions to facilitate spike conduction in Ia afferents, as well as directly evoke spikes in Ia afferents. This dorsal population is also recurrently activated by Ia afferents in a classic trisynaptic loop, where Ia or cutaneous afferents activate first order neurons (Lin *et al*., 2023) that in turn widely activate these dorsal GAD2 neurons that innervate GABA_A_ receptors on Ia afferents to cause PAD and associated spikes that again further activate this GAD2 pathway. This serves to both facilitate conduction in Ia afferents and amplify spikes, but is readily mistaken for presynaptic inhibition because it leads to post-activation depression of Ia synapses that are repeatedly activated.

We also demonstrate that following chronic SCI, the number of Ia terminal GABA_B_ receptors and their hyperpolarizing effect on the Ia afferent are markedly reduced. In mice this is associated with reduced GABA_B_ receptor-mediated post-activation depression of Ia-EPSPs within the long time-window of the GABA_B_ receptor effect (∼ 1 second). In human participants, post-activation depression of H-reflexes, induced either from repetitive activation of the same Ia afferent or from conditioning of an antagonist nerve, is also reduced after chronic SCI, in the same long time-window, consistent with a loss of GABA_B_ receptors in humans. Indeed, SCI participants taking the GABA_B_ receptor agonist baclofen exhibited increased post-activation depression of H-reflexes within the estimated time-window of the GABA_B_ receptor effect (∼ 1 second). The observed marked reduction in Ia afferent terminal GABA_B_ receptors and their action after SCI suggests that with repeated sensory activation there may be increases in the activation of VCa^2+^ channels that speed up vesicle replenishment and neurotransmitter release to reduce post-activation depression and contribute to the exaggerated reflex activation of spinal motoneurons and interneurons in spasticity. Whether similar changes in axonal GABA_B_ receptor expression occur elsewhere in the spinal cord in SCI is unknown, though worth examining, such as for interneurons (Delgado-Ramirez *et al*., 2023) and descending axon terminals (Jimenez *et al*., 1991) innervated by GABAergic neurons .

### Changes in spinal GABA networks after SCI

Following SCI there are complex changes in spinal GABA neurons below the injury due to their dependence on sensorimotor activity (Mende *et al*., 2016; Lalonde & Bui, 2021). Although the number of GABA (GAD2) interneurons with axo-axonic connections to Ia afferents that release synaptic GABA does not change after SCI, the number of GAD2 boutons that innervate the Ia terminals decrease (Kapitza *et al*., 2012; Smith *et al*., 2017; Khalki *et al*., 2018), potentially from a reduction in the activity-dependent release of BDNF and glutamate from sensory afferents (Mende *et al*., 2016; Lalonde & Bui, 2021). The reduced number of terminal GABAergic boutons after SCI might exacerbate the action of the decreased GABA_B_ receptors on the Ia afferent terminal found here, with both combining to markedly reduce presynaptic inhibition of Ia afferents and subsequent post-activation depression of the Ia-EPSP as detailed below.

#### Mechanisms of GABA_B_ receptor-mediated presynaptic inhibition

Previously we have proposed that presynaptic inhibition during repetitive activation of Ia-EPSPs (i.e., homosynaptic post-activation depression) is produced by collaterals of the Ia afferent that activate a recurrent GABAergic (GAD2) pathway with axo-axonic connections back onto the same Ia terminal that, in turn, synapse onto GABA_B_ receptors [Abstract figure and (Metz *et al*., 2023a). The presence of this circuit is supported here by the confirmation of GABA_B_ receptors being mainly located on the Ia terminal in close juxtaposition to GAD2-positive terminal boutons and the finding that repetitive optogenetic activation of ventral GAD2//ChR2 neurons mimics the post-activation depression produced by repetitive Ia activation, with both being strongly reduced by GABA_B_ receptor blockade. The selective application of light to the dorsal and ventral spinal cord further suggests the presence of two populations of GAD2 neurons detailed above, one with projections primarily to dorsally located Ia afferent nodes that produces PAD (via GABA_A_ receptors) and the other with primarily ventral projections onto the Ia terminal and motoneuron that initiates Ia presynaptic inhibition (via GABA_B_ receptors) and postsynaptic motoneuron inhibition (via GABA_A/B_ receptors), respectively. This was revealed by the dorsal light producing mainly PAD in Ia afferents and facilitation of Ia-EPSPs when evoked within the PAD window, whereas the ventral light not producing PAD and instead, producing an early IPSP in the motoneurons that was then followed by a long-latency suppression of the Ia-EPSP after the membrane potential returned to baseline. Thus, during repeated activation of Ia afferents, it is likely that both groups of GAD2 neurons are activated to increase conduction in Ia afferents (from dorsal activation of PAD) and at the same time, activate the recurrent GAD2-GABA_B_ pathway to produce long-lasting, post-activation depression of the Ia-EPSP. Moreover, if the sensory stimulation is strong enough, the resulting PAD may evoke orthodromic spikes in the Ia afferent that produce post-activation depression of subsequent Ia-EPSPs. Here post-activation depression is produced when the PAD spikes, mediated by GABA_A_ receptors, evoke a conditioning Ia-EPSP in addition to recruiting the recurrent GAD2-GABA_B_ receptor pathway, so a later evoked test Ia-EPSP becomes suppressed. Such a mechanism may explain why the application of the GABA_A_ receptor antagonists L655,708 also reduces post-activation depression (Lee-Kubli & Calcutt, 2014; Hernandez-Reyes *et al*., 2019), potentially by reducing PAD-evoked spikes and their subsequent conditioning of the Ia-EPSP and activation of the recurrent GAD2-GABA_B_ receptor pathway.

Once activated, there are two possible mechanisms whereby GABA_B_ receptors located on the terminals of Ia afferents can produce presynaptic inhibition. First, GABA_B_ receptors can facilitate GIRK channels and the outflow of potassium ions to hyperpolarize the afferent membrane potential (Luscher & Slesinger, 2010). This was observed by the hyperpolarization of the membrane potential of the Ia afferent during application of the GABA_B_ receptor agonist baclofen (Fig. 5). However, it is important to note that the membrane potential recordings were made with grease gap connections to the cut proximal dorsal root, far from the Ia afferent terminal, explaining the very small amplitude signals (<0.5 mV) that were measured. Thus, intracellular recordings closer to the Ia afferent terminal [as done in (Lucas-Osma *et al*., 2018) are needed to verify these results. If the amount of hyperpolarization of the Ia afferent terminal by GABA_B_ receptors is indeed very small, it is likely not the main driver of presynaptic inhibition as this would produce a small reduction in the size of the incoming action potential and calcium inflow to the presynaptic Ia terminal (Hari *et al*., 2022).

The second mechanism by which GABA_B_ receptors produce presynaptic inhibition of the Ia terminal is through its inhibition of VCa^2+^ channels to reduce calcium entry and vesicle release from the presynaptic terminal following the arrival of an action potential (Dolphin & Scott, 1986; Lev-Tov *et al*., 1988; Curtis *et al*., 1997; Abe *et al*., 2019). The finding that the GABA_B_ receptor agonist baclofen, at doses that do not affect the VCa^2+^ channels on the motoneuron, reduces the Ia-EPSP supports the mechanism of GABA_B_ receptors reducing calcium entry and neurotransmitter release in the Ia terminal (Lev-Tov *et al*., 1988; Li *et al*., 2004b). GABA_B_ receptors also facilitate post-activation depression during repeated activation of the same Ia-EPSP [see also (Fink, 2013)], potentially by reducing the replenishment rate of neurotransmitter vesicles. As outlined in the Introduction, Ca^2+^-calmodulin in the presynaptic terminal of the Calyx of Held increases the replenishment rate and the readily releasable pool of neurotransmitters via the Munc13-1 pathway (Sakaba & Neher, 2001a; Lipstein *et al*., 2013). In support of this, the facilitation of GABA_B_ receptors with baclofen, and its inhibition of terminal VCa^2+^ channels and calcium calmodulin, reduces vesicle recovery during repetitive activation of the Calyx of Held synapse to increase post-activation depression (Sakaba & Neher, 2003). A similar mechanism could explain the facilitation of post-activation depression by baclofen during repetitive activation of the Ia-motoneuron synapse during H-reflex activation.

The reduction in vesicle replenishment from GABA_B_ receptor activation appears to be long-lasting. For example, blockade of GABA_B_ receptors with CGP55845 in intact mice reduced post-activation depression of Ia-EPSPs for up to 1 second and this might be attributed to a prolonged G-protein coupled response and its effect on the terminal VCa^2+^ channels. Although this pharmacological result is compelling, direct recordings with Ca^2+^ imaging of the Ia terminal are needed to verify the effect of terminal GABA_B_ receptors on presynaptic VCa^2+^ channel activation and subsequent neurotransmitter release and vesicle replenishment (Kavalali & Jorgensen, 2014). In fact, the process of presynaptic inhibition, defined as a decrease in neurotransmitter release at a presynaptic terminal (Quevedo, 2009), remains to be directly measured in the Ia-motoneuron synapse.

#### Historical confusion of PAD with presynaptic inhibition

Historically, GABA_A_ receptor-mediated PAD has been confused with GABA_B_ receptor-mediated presynaptic inhibition in the Ia afferent because PAD is so excitatory that it readily evokes spikes in Ia afferents that consistently travel orthodromically to evoke EPSPs in motoneurons, leading to post-activation depression, as we detail here. Futhermore, while these spikes have a lower probability of travelling antidromically out the dorsal root, they do so in both in vivo and in vitro preparations including during locomotion, demonstrating the univeral importance of PAD-evoked spikes (Eccles *et al*., 1962; Gossard & Rossignol, 1990; Lucas-Osma *et al*., 2018). The indirect monosynaptic excitation of Ia-EPSPs via PAD-evoked spikes leads to post activation depression (GABA_B_ and transmitter depletion mediated) of subsequently tested monosynaptic EPSPs and H-reflexes that outlasts PAD by hundreds of ms, making it unlikely from the outset that PAD directly caused presynaptic inhibition (Curtis & Eccles, 1960; Hari *et al*., 2022; Metz *et al*., 2023a). This excitatory action of PAD and subsequent paradoxical post-activation depression is readily mistaken for presynaptic inhibition, because it depends on PAD that is GABA_A_ mediated, and so is sensitive to GABA_A_ blockers applied even locally the spinal cord as mentioned above [see also (Stuart & Redman, 1992; Hari *et al*., 2022)]. Although largely ignored, such PAD-related EPSPs are evident in the original Frank and Fuortes publications that led to the modern concept of presynaptic inhibition [see Fig. 6 in (Frank, 1959)] and appear in many other publications (Duchen, 1986), including in human near-nerve recordings (Shefner *et al*., 1992), in the work of Redman involving in vivo GABA_A_ receptor antagonists (Stuart & Redman, 1992), and the in vitro work from the Jessel laboratory (Fink, 2013). To further complicate matters, these PAD-evoked EPSPs can be masked by IPSPs that invariably are activated by GAD2 neurons which release GABA onto motoneurons and simultaneously to afferents terminals (in a triadic arrangement), as we show here, though this does not affect afferent terminal post-activation depression (Stuart & Redman, 1992; Pierce & Mendell, 1993; Hari *et al*., 2022). Importantly, while phasic PAD and associated PAD-evoked spikes (DRRs) do not change much with SCI (Lin *et al*., 2023), PAD-evoked spikes have a much smaller inhibitory action on subsequently tested EPSPs (Fig. 1J), which is accounted for by reductions in afferent GABA_B_ receptors with SCI, and provides more support for the accumulating evidence that PAD is not generally associated with presynaptic inhibition in proprioceptors.

#### Loss of terminal GABA_B_ receptors and its effect on post-activation depression

In mice, a reduced number of GABA_B_ receptors on the Ia terminal after SCI is associated with a reduction in post-activation depression of Ia-EPSPs, mediated either from direct Ia activation (Fig. 4) or from PAD-evoked spikes (Fig. 1I). Interestingly in intact mice, it was possible to reduce post-activation depression to levels observed in chronically injured mice by blocking GABA_B_ receptors with CGP55845, specifically at ISIs up to 1 second within the time window of the GABA_B_ receptor effect. As described above, a reduction in terminal GABA_B_ receptors shown here, combined with a reduction in GAD2 bouton innervation (Kapitza *et al*., 2012; Smith *et al*., 2017; Khalki *et al*., 2018), would lead to a reduced inhibition of VCa^2+^ channels and more calcium inflow into the presynaptic terminal. A higher amount of intracellular Ca^2+^ could in theory activate more calmodulin to increase the replenishment rate of vesicles so that there would be a larger, readily releasable pool of neurotransmitters to reduce post-activation depression (Sakaba & Neher, 2001a). The 50% reduction in the number of terminal GABA_B_ receptors after chronic SCI also reduced, by the same amount, the hyperpolarization of the mean membrane potential following application of the GABA_B_ receptor agonist baclofen, likely by reducing the activation of GIRK channels and the outward flow of K^+^ ions. The reduced efficacy of baclofen in activating GABA_B_ receptors following spinal cord injury is also demonstrated by the doubling of the half-maximal effective concentration (EC_50_) of baclofen needed to reduce the Ia-EPSP (Li *et al*., 2004b). Thus in rodents, an increased activation of VCa^2+^ channels on the Ia terminal and vesicle release after chronic SCI likely allows for larger Ia evoked EPSPs (Li *et al*., 2004a) and reduced post-activation depression that may contribute to hyperactive stretch reflexes.

Similar to mice, post-activation depression of the largely Ia-mediated soleus H-reflex from repeated TN stimulation, or from a conditioning antagonist CPN stimulation, was reduced in human participants with chronic spinal cord injury compared to intact controls, especially in participants who were not taking the GABA_B_ receptor agonist baclofen as a daily antispastic treatment (Fig. 10). The difference in post-activation depression of H-reflexes between injured participants taking vs not taking baclofen lasted out to 2 seconds compared to only 1 second for Ia-EPSPs during modulation of GABA_B_ receptors in mice, potentially because the initiation of action potentials evoked during the H-reflex is more sensitive to changes in presynaptic inhibition.

In human participants, the profile of post-activation depression of the soleus H-reflex during conditioning by antagonist CPN stimulation was similar to that produced by direct activation of the soleus Ia afferents from TN stimulation. As shown in mice, such H-reflex suppression could have been mediated by the CPN afferents producing an orthodromic, PAD-evoked spike in the soleus Ia afferents that then evoked a conditioned Ia-EPSP to then suppress H-reflexes evoked many seconds later (Metz *et al*., 2023a). The activation of an early soleus response from the CPN stimulation is an indication that the PAD-evoked spikes reached the Ia terminal and activated the soleus motoneurons. Despite a larger early soleus response to CPN stimulation, the amount of post-activation depression of H-reflexes in participants with SCI was less compared to controls. This could result from a facilitated rate of recovery of vesicle replenishment after SCI due to tonically increased levels of presynaptic calcium (Deng *et al*., 2019). In addition, the PAD-evoked spikes in the soleus Ia afferents may also activate a weakened recurrent GAD2-GABA_B_ receptor pathway due to the reduced number of GAD2 boutons and GABA_B_ receptors on the Ia terminal. In intact cats, suppression of the extensor soleus/gastrocnemius Ia-EPSP from a prior conditioning stimulation of the flexor posterior biceps-semitendinosus (PBST) nerve is reduced by the intravenous administration of the GABA_B_ receptor antagonist CGP46381 (Curtis & Lacey, 1998). The duration of the reduced Ia-EPSP suppression occurred at ISIs between 40 and 800 ms, again within the duration of the GABA_B_ receptor effect. Likewise, we show here that after SCI where the number of GABA_B_ receptors is greatly reduced, the amount of Ia-EPSP suppression following PAD-evoked spikes from dorsal light application is greatly reduced (Fig. 1J). Collectively, these findings suggest that after SCI, antagonist afferents activate a weakened recurrent GAD2-GABA_B_ receptor pathway via PAD evoked spikes in the test Ia afferent that results in a decrease of post-activation depression.

### Functional role of post-activation depression

A decrease of post-activation depression of Ia-EPSPs might contribute to the hyperreflexia and clonus after SCI, especially during repetitive activation of Ia afferents during cyclical movements such as walking. Like electrical stimulation, synchronous activation of Ia afferents during a muscle stretch, tendon tap or isometric contraction can also produce subsequent post-activation depression of H-reflexes, which are also reduced after SCI (Crone & Nielsen, 1989; Nielsen *et al*., 1993; Grey *et al*., 2008). Thus, post-activation depression of Ia-EPSPs is likely a physiological process that controls reflex activation of spinal motoneurons during natural motor behaviour. Interestingly, post-activation depression is greatly reduced during a superimposed voluntary contraction (Stein *et al*., 2007; Clair *et al*., 2011; Ozyurt *et al*., 2020), potentially from postsynaptic facilitation of the motoneuron and/or by descending inputs reducing the activation of GABA_B_ receptors on the Ia terminal. Whether residual voluntary contractions or involuntary activity during spasms affects post-activation depression after SCI is unknown. However, we observed that during a buildup of background EMG with faster stimulation rates (e.g., 500 ms ISIs), suppression of H-reflexes was reduced, and these trials were removed from analysis. In summary, the decrease in neurotransmitter release and increase in post-activation depression by facilitating GABA_B_ receptors on the Ia terminal with baclofen likely explains the antispastic effect of this drug in SCI. Although other neurological disorders also show reduced post-activation depression of H-reflexes, such as MS (Leon *et al*., 2023), ALS (Zhou *et al*., 2022), stroke (Milanov, 1992; Faist *et al*., 1994; Aymard *et al*., 2000), brain injury (Koelman *et al*., 1993) and cerebral palsy (Achache *et al*., 2010), baclofen might not be as effective compared to SCI given the greater amounts of involuntary descending inputs that also increase the excitability spinal circuits in these disorders (Mottram *et al*., 2009). In summary, these findings reveal the important role of GABA_B_ receptors on the Ia afferent terminal in presynaptic inhibition and why post-activation depression may be reduced after chronic SCI to help understand the potential mechanisms underlying hyperreflexia in spasticity.

## Acknowledgments

We thank Ms Jennifer Duchcherer and Leo Sanelli for technical assistance.

## Data Availability

Data can be provided upon request and statistical results are summarized in the Supplemental Tables document.

## Additional Information

### Author contributions

K.M., K.H., A.L.O., D.J.B. and M.A.G. conceived and designed research; K.M., K.H., A.L.O., R.M., I.C.M., S.Y., J.F.Y., D.J.B. and M.A.G. performed experiments; K.M., K.H., A.L.O., R.M., T.O.A., I.C.M., S.Y., J.F.Y., D.J.B. and M.A.G. analyzed the data and interpreted the results of experiments; K.M., K.H., A.L.O., D.J.B. and M.A.G. prepared figures; K.M., K.H., D.J.B. and M.A.G. drafted and revised the manuscript and all authors edited the manuscript.

All authors approved the final version of the article and agreed to be accountable for all aspects of the work. The authors confirm that all persons designated as authors are qualified.

