## Supplemental Tables for "GABA_B_ receptors mediate presynaptic inhibition of proprioceptive afferents and have reduced action after spinal cord injury" (from another version of this preprint: version 1, 2026-01-22)

| ID | Age | Sex | Years | Injury | Leg move | CPN-TN | TN-TN |
|---|---|---|---|---|---|---|---|
| 1 | 25 | M | .5 | T5/6 | No | ✓ (blue) | ✓ (blue) |
| 2 | 31 | M | 11 | C5 | Yes | ✓ (blue) | ✓ (blue) |
| 3 | 23 | M | 6 | C2/3 | No | ✓ (blue) | ✓ (blue) |
| 4 | 61 | F | 3 | C4/5 | Yes | ✓ (blue) | ✓ (blue) |
| 5 | 31 | M | 5.5 | T6 | No | ✓ (blue) | ✓ (blue) |
| 6 | 58 | M | 3.5 | C3/4 | No | ✓ (red) | ✓ (red) |
| 7 | 20 | M | 3.5 | T8 | No | ✓ (red) | ✓ (red) |
| 8 | 65 | F | 6.5 | T12 | Yes | ✓ (red) | ✓ (red) |
| 9 | 28 | M | 15 | C4/5 | Yes | ✓ (red) | ✓ (red) |
| 10 | 25 | M | 1.5 | T5 | Yes | ✓ (red) | ✓ (red) |
| 11 | 36 | M | 11.5 | C4 | No | ✓ (red) | ✓ (red) |
| 12 | 26 | F | 10 | C4/5 | No | ✓ (red) | ✓ (red) |
| 13 | 24 | M | 4 | C5 | No | ✓ (red) | ✓ (red) |
| 14 | 24 | F | 3 | T5 | No | ✓ | ✓ |
| 15 | 29 | F | 11 | C4/5/6 | No | ✓ | ✓ |
| 16 | 29 | M | 12 | T4 | Yes | ✓ | ✓ |
| 17 | 35 | M | 3 | T7 | No | ✓ (blue) | ✓ |
| 18 | 51 | M | 13.5 | T12 | Yes | ✓ (blue) | ✓ (blue) |
| 19 | 33 | M | 5.5 | T2 | No | ✓ | ✓ |
| 20 | 21 | M | 4 | C4 | No | ✓ | ✓ |
| 21 | 57 | M | 3.5 | T2 | No | ✓ | ✓ |
| 22 | 64 | M | .5 | C4 | Yes | ✓ (blue) | |
| 23 | 21 | F | 5 | T6 | No | ✓ (red) | |
| 24 | 28 | M | 4.5 | T4 | Yes | | ✓ |
| 25 | 45 | M | 7 | C6 | Yes | | ✓ |
| 26 | 54 | F | 11 | T2 | Yes | | ✓ |
| 27 | 67 | M | 24.5 | C4 | Yes | | ✓ |
| 28 | 64 | M | 33.5 | T12 | Yes | | ✓ |
| 29 | 47 | M | 11 | T12 | Yes | | ✓ |
| 30 | 55 | M | 18 | C6-7 | Yes | | ✓ |
| 31 | 52 | M | 1.5 | T10 | Yes | | ✓ |
| 32 | 47 | M | 1 | T12 | Yes | | ✓ |
| 33 | 64 | M | 1 | C3 | Yes | | ✓ |
| 34 | 57 | F | 10.5 | C6 | Yes | | ✓ |
| 35 | 30 | F | 4.2 | C6 | Yes | | ✓ |
| 36 | 44 | F | 22 | T6/7 | No | | ✓ |
| **36** | **40.6±15.7** | **26 M** | **8.1±7.3** | **C2-T12** | **16N/20Y** | **23** | **34** |

**Supplemental Table 1. Participant demographics, injury and experiment details**. Age, sex, number of years post-injury, neurologic level of injury and whether a participant could voluntarily activate one or more muscles in their leg (Yes/No). Checkmarks indicate if the participant participated in the CPN or RDD experiment. ✓ (red) = not on baclofen, ✓ (blue) = on baclofen whose data were used for comparisons in Figure 10.

| **A) Figure 1D** | **% change EPSP** | | **Test value** | **P-value** | **Statistic** |
|---|---|---|---|---|---|
| *Ventral light* | -64.903 (13.086) | | | | |
| Ventral light + CGP | -11.757 (12.571) | | t = -9.957 | **< 0.001** | Paired t |
| Ventral light in Gad2- | -3.964 (4.827) | | t = -11.604 | **< 0.001** | Unpaired t |
| **Figure 1E** | **IPSP Amplitude (mV)** | | **Test value** | **P-value** | **Statistic** |
| *Ventral light* | -0.0589 (0.0254) | | | | |
| Ventral light + Gbz | -0.0116 (0.0112) | | t = -5.026 | **< 0.001** | Paired t |
| Dorsal light | 0.0011 (0.0010) | | T = 54.000 | **0.005** | MW |
| Dorsal light + Gbz | 0.0030 (0.0035) | | T = 54.000 | **0.005** | MW |
| **Figure 1F** | **PAD Amplitude (mv)** | | **Test value** | **P-value** | **Statistic** |
| Ventral light | 0.0282 (0.0246) | | T = 47.000 | **<0.001** | MW |
| Ventral light + Gbz | 0.0000 (0.0000) | | T = 45.000 | **<0.001** | MW |
| *Dorsal light* | 0.486 (0.358) | | | | |
| Dorsal light + Gbz | -0.0006 (0.0055) | | t = 4.347 | **0.002** | Paired t |
| **Figure 1H** | **% change EPSP with dorsal light** | | **Test value** | **P-value** | **Statistic** |
| *During PAD* | 64.914 (48.097) | | | | |
| During PAD + CGP | 163.914 (122.201) | | t = -2.450 | **0.050** | Paired t |
| During PAD+CGP+Gbz | -4.367 (14.822) | | t = 3.156 | **0.020** | Paired t |
| After PAD | -9.5124 (8.670) | | t = 4.425 | **0.004** | Paired t |
| **Figure 1J** | **Difference in % change in EPSP** | | **Test value** | **P-value** | **Statistic** |
| Doral light control | -50.8001 (39.838) | | T = 84.000 | **<0.001** | MW |
| Dorsal light SCI | -5.617 (48.432) | | T = 63.000 | 0.463 | MW |
| Ventral light control | -4.143 (16.899) | | t = 0.649 | 0.529 | Unpaired t |
| **B) Figure 2C** | **GAD2/A3** | **GAD2/A3+light** | **Test value** | **P-value** | **Statistic** |
| % change EPSP | -59.283(11.850) | -31.958 (19.135) | t = - 4.440 | **0.003** | Paired t |
| | **WT** | **WT+CGP55** | | | |
| % change EPSP | -58.234(11.862) | -34.094 (11.068) | t = - 4.089 | **0.006** | Paired t |
| **Figure 2G** | **% change EPSP** | | **Test value** | **P-value** | **Statistic** |
| *Ventral light* | -63.250 (16.264) | | | | |
| Ventral light + CGP | -10.000 (11.071) | | t = - 10.270 | **< 0.001** | Paired t |
| *DR train* | -58.039 (8.130) | | | | |
| DR train + CGP | -34.075 (14.177) | | t = - 3.870 | **0.008** | Paired t |
| *Vent light+DR train* | -21.304 (6.946) | | | | |
| Vent light+DR+CGP | -7.912 (6.732) | | t = - 2.872 | **0.028** | Paired t |
| **Figure 2H** | **Changed in Vm (mV)** | | **Test value** | **P-value** | **Statistic** |
| Repeated sensory | -0.0135 (0.004) | | T = 28.000 | **< 0.001** | MW |
| Repeated sensory +CGP | -0.0141 (0.006) | | T = 52.500 | **0.001** | MW |
| Repeated sensory + Gbz | -0.0311 (0.018) | | t = -3.089 | **0.037** | Unpaired t |
| Repeated light | 0.00974 (0.007) | | T = 77.000 | **< 0.001** | MW |
| Repeated light + CGP | 0.00514 (0.008) | | T = 63.000 | 0.209 | MW |
| Repeated light + Gbz | 0.0023 (0.004) | | t = 0.971 | 0.387 | Unpaired t |
| **C) Figure 3** | **Control (%)** | **SCI (%)** | **Power** | **P-value** | **Statistic** |
| GABAbr / terminal | 100 | 58.530 (18.600) | T = 14.000 | **0.008** | MW |
| GABAbr / MN | 100 | 113.939(23.738) | T = 30.000 | 0.690 | MW |
| GABAbr terminal/MN | 49.109 (9.084) | 25.652 (6.820) | t = 4.617 | **0.002** | Unpaired t |
| Terminals w receptors | 93.416 (2.861) | 91.038 (4.843) | t = 0.945 | 0.372 | Unpaired t |
| **D) Figure 5** | **Control** | **SCI** | **Power** | **P-value** | **Statistic** |
| Decrease Vm (mV) | -0.271 ±0.110 | -0.120 ± 0.050 | t = - 2.212 | **0.044** | Unpaired t |
| **E) Figure 6** | **Control** | **SCI** | **Power** | **P-value** | **Statistic** |
| Hmax (mV) | 3.577 (1.621) | 2.502 (1.699) | T = 706.500 | **0.023** | MW |
| Mmax (mV) | 7.557 (3.298) | 4.984 (2.766) | T = 697.000 | **0.004** | MW |
| Hmax/Mmax (x100%) | 50.535 (16.381) | 51.702 (21.769) | t = 0.207 | 0.837 | Unpaired t |

**Supplemental Table 2.** Mean and standard deviation (SD) of variables presented in the bar graphs of Figures 1, 2, 3, 5 and 6 along with results of statistical comparisons for unpaired and paired Student's t-tests or Mann-Whitney Rank Sum test (MW). Variables that are compared to are marked with italics.

**A) Fig. 1D** % change in Ia-EPSP from prior ventral light stimulation in GAD2+//ChR2 mice before and after CPG55845 and in GAD2- mice with paired and unpaired Student's t-test comparisons to ventral light only; **Fig. 1E&F** amplitude of ventral and dorsal light evoked IPSP (E) and PAD (F) before and after gabazine (Gbz) with paired comparison to ventral (E) or dorsal (F) light only; **Fig. 1H** % change in EPSP during PAD without and with CPG55845 or CGP+Gbz and after PAD with comparison to EPSP during PAD; and **Fig. 1J** difference in % change of EPSP following PAD-evoked spikes from dorsal light in control and chronic SCI mice and from non-spiking ventral light, all variables in comparison to 0% change.
**B) Fig. 2C** % change of 4$^{th}$ EPSP compared to 1$^{st}$ EPSP during repetitive DR stimulation in GAD2//Arch3+ mice before and during suppression of GAD2 neurons by light and before and after CPG55845 application in wild-type (WT) mice, results of paired comparisons; **Fig. 2G** % change in 4$^{th}$ EPSP vs 1$^{st}$ EPSP during repetitive ventral light train, DR stimulation train or both with and without CGP55845, all paired comparisons; **Fig. 2H** Change in motoneuron potential following repeated DR or ventral light stimulation with and without CGP845 and Gbz, unpaired comparisons to 0 mV change.
**C) Fig. 3B** mean intensity of $GABA_B$ receptors on Ia terminals and motoneurons in SCI mice as a percentage of control mice, unpaired comparisons; Fig. 3C mean intensity of $GABA_B$ receptors on Ia terminals relative to the intensity on motoneurons, unpaired comparison; Fig. 3D number of Ia terminals with $GABA_B$ receptors as a percentage of total labeled terminals, unpaired comparison.
**D) Fig. 5** mean (SD) decrease in baseline DC membrane potential (Vm) during application of baclofen, unpaired comparisons between control mice and mice with chronic SCI. **E)** Mean (SD) $GABA_B$ receptor density over: Ia terminal in mice with chronic SCI as a % of intensity over Ia terminal in intact mice; motoneuron in SCI mice as % of motoneuron in intact mice and Ia terminal as % in respective motoneuron of SCI and intact mice.
**E) Fig. 6B** mean (SD) of Hmax, Mmax and Hmax/Mmax ratio in participants with and without SCI (control) and results of unpaired comparisons.

| **Figure 4: RDD of Ia EPSPs in mice** | | | | |
|---|---|---|---|---|
| **A: WT vs SCI** | **DF** | **Test value** | **P value** | **Test** |
| Group | 1 | F = 20.081 | **P < 0.001** | Two-way ANOVA |
| ISI | 3 | F = 64.044 | **P < 0.001** | Two-way ANOVA |
| Group x ISI | 3 | F = 12.477 | **P < 0.001** | Two-way ANOVA |
| 0.5s ISI | | t = 7.638 | **P < 0.001** | Holm-Sidak |
| 1s ISI | | t = 2.906 | **P = 0.005** | Holm-Sidak |
| **B: WT vs WT CGP** | | | | |
| Group | 1 | F = 21.649 | **P < 0.001** | Two-way ANOVA |
| ISI | 3 | F = 79.900 | **P < 0.001** | Two-way ANOVA |
| Group x ISI | 3 | F = 5.417 | **P = 0.002** | Two-way ANOVA |
| 0.5 s ISI | | t = 5.021 | **P < 0.001** | Holm-Sidak |
| 1 s ISI | | t = 4.008 | **P < 0.001** | Holm-Sidak |
| **C: SCI vs SCI CGP** | | | | |
| Group | 1 | F = 0.433 | P = 0.512 | Two-way ANOVA |
| ISI | 3 | F = 49.987 | **P < 0.001** | Two-way ANOVA |
| Group x ISI | 3 | F = 5.990 | **P = 0.001** | Two-way ANOVA |
| 0.5 s ISI | | t = 3.085 | **P = 0.003** | Holm-Sidak |
| 1 s ISI | | t = 2.886 | **P = 0.005** | Holm-Sidak |

**Supplemental Table 3.** Fig. 4 Results of two-way ANOVA comparing % change of Ia-EPSP with main factors "group" and "ISI" between: **A)** control and chronic SCI mice, **B)** control mice before and during application of CGP55845 and **C)** SCI mice before and during application of CGP55845.

| **Figure 7: CPN conditioning of soleus H-reflex at short ISIs in humans.** | | | | |
|---|---|---|---|---|
| **Controls: H-reflexes** | **DF** | **Test value** | **P value** | **Test** |
| 1.0 x MT | 5 | H = 15.094 | **P = 0.010** | One-Way ANOVA on Ranks |
| Diff from 0% change | 6 | H = 21.116 | **P = 0.002** | One-way ANOVA on Ranks |
| 30 ms ISI | | **q = 5.418** | **P < 0.05** | Tukey |
| 1.5 x MT | 5 | H = 24.537 | **P < 0.001** | One-Way ANOVA on Ranks |
| Diff from 0% change | 6 | H = 34.952 | **P < 0.001** | One-way ANOVA on Ranks |
| 15 ms ISI | | **q = 4.799** | **P < 0.05** | Tukey |
| 100 ms ISI | | **q = 6.636** | **P < 0.05** | Tukey |
| **SCI: H-reflexes** | | | | |
| 1.0 x MT | 5 | F = 1.671 | P = 0.146 | One-Way ANOVA |
| 1.5 x MT | 5 | H = 27.023 | **P < 0.001** | One-Way ANOVA on Ranks |
| Diff from 0% change | 6 | H = 36.065 | **P < 0.001** | One-way ANOVA on Ranks |
| 15 ms ISI | | **Q = 3.225** | **P < 0.05** | Dunn's Method |
| 200 ms ISI | | **Q = 3.837** | **P < 0.05** | Dunn's Method |
| **SCI vs Control** | | | | |
| 1.0 x MT Group | 1 | F = 6.478 | **P = 0.012** | Two-Way ANOVA |
| 1.0 x MT ISI | 5 | F = 3.127 | **P = 0.010** | Two-Way ANOVA |
| 1.0 x MT Group x ISI | 5 | F = 2.131 | P = 0.063 | Two-Way ANOVA |
| 30 ms ISI | | t = 4.827 | **P < 0.001** | Student's t-test, Bonferroni |
| 1.5 x MT Group | 1 | F = 11.982 | **P < 0.001** | Two-way ANOVA |
| 1.5 x MT ISI | 5 | F = 5.519 | **P < 0.001** | Two-Way ANOVA |
| 1.5 x MT Group x ISI | 5 | F = 1.893 | P = 0.098 | Two-Way ANOVA |
| 100 ms ISI | | T = 140.000 | **P = 0.005** | Mann-Whitney, Bonferroni |
| **EMG (30 to 80 ms)** | | | | |
| 1.0 x MT Group | 1 | F = 0.0159 | P = 0.900 | Two-Way ANOVA |
| 1.0 x MT bin | 5 | F = 6.883 | **P < 0.001** | Two-Way ANOVA |
| 1.0 x MT Group x ISI | 5 | F = 1.483 | P = 0.197 | Two-Way ANOVA |
| 1.5 x MT Group | 1 | F = 8.731 | **P = 0.004** | Two-Way ANOVA |
| 1.5 x MT bin | 5 | F = 15.785 | **P < 0.001** | Two-Way ANOVA |
| 1.5 x MT Group x ISI | 5 | F = 1.317 | P = 0.258 | Two-Way ANOVA |

**Supplemental Table 4.** Fig. 7 Results of one and two-way ANOVAs comparing % change of H-reflex and early soleus response across short ISIs (3 to 200 ms) and between SCI and control groups for the 1.0 x MT and 1.5 x MT conditioning CPN stimulation intensities.

| **Figure 9: CPN-TN and TN-TN conditioning at long ISIs in humans** | | | | |
|---|---|---|---|---|
| **Control: CPN-TN** | **DF** | **Test value** | **P value** | **Test** |
| Diff from 0% change | 5 | F = 23.151 | **P < 0.001** | One-way ANOVA |
| 500 ms ISI | | t = 9.780 | **P < 0.001** | Holm-Sidak |
| 1000 ms ISI | | t = 7.088 | **P < 0.001** | Holm-Sidak |
| 2000 ms ISI | | t = 3.927 | **P < 0.001** | Holm-Sidak |
| 2500 ms ISI | | t = 2.912 | **P = 0.005** | Holm-Sidak |
| **SCI: CPN-TN** | | | | |
| Diff from 0% change | 5 | H = 35.637 | **P < 0.001** | One-Way ANOVA on Ranks |
| 500 ms ISI | | Q = 4.276 | **P < 0.05** | Dunn's Method |
| 1000 ms ISI | | Q = 4.955 | **P < 0.05** | Dunn's Method |
| 2000 ms ISI | | Q = 4.762 | **P < 0.05** | Dunn's Method |
| 2500 ms ISI | | Q = 2.377 | P > 0.05 | Dunn's Method |
| **SCI vs Control CPN-TN** | | | | |
| Group | 1 | F = 22.658 | **P < 0.001** | Two-Way ANOVA |
| ISI | 4 | F = 9.728 | **P < 0.001** | Two-Way ANOVA |
| Group x ISI | 4 | F = 3.422 | **P = 0.010** | Two-Way ANOVA |
| 500 ms ISI | | t = 5.273 | **P < 0.001** | Holm-Sidak |
| 1000 ms ISI | | t = 2.329 | **P = 0.021** | Holm-Sidak |
| **Control: TN-TN** | | | | |
| Diff from 0% change | 4 | H = 64.679 | **P < 0.001** | One-way ANOVA |
| 500 ms ISI | | q = 9.672 | **P < 0.05** | Tukey |
| 1000 ms ISI | | q = 8.090 | **P < 0.05** | Tukey |
| 2000 ms ISI | | q = 5.766 | **P < 0.05** | Tukey |
| 5000 ms ISI | | q = 2.292 | P > 0.05 | Tukey |
| **SCI: TN-TN** | | | | |
| Diff from 0% change | 4 | H = 79.815 | **P < 0.001** | One-way ANOVA |
| 500 ms ISI | | Q = 5.686 | **P < 0.05** | Dunn's Method |
| 1000 ms ISI | | Q = 8.022 | **P < 0.05** | Dunn's Method |
| 2000 ms ISI | | Q = 5.770 | **P < 0.05** | Dunn's Method |
| 5000 ms ISI | | Q = 3.270 | **P < 0.05** | Dunn's Method |
| **SCI vs Control: TN-TN** | | | | |
| Group | 1 | F = 35.380 | **P < 0.001** | Two-Way ANOVA |
| ISI | 3 | F = 39.767 | **P < 0.001** | Two-Way ANOVA |
| Group x ISI | 3 | F = 3.516 | **P = 0.017** | Two-Way ANOVA |
| 500 ms ISI | | t = 4.951 | **P < 0.001** | Holm-Sidak |
| 1000 ms ISI | | t = 3.056 | **P = 0.003** | Holm-Sidak |
| 2000 ms ISI | | t = 2.752 | **P = 0.007** | Holm-Sidak |
| **Control CPN-TN vs TN-TN** | | | | |
| Group | 1 | F = 12.543 | **P < 0.001** | Two-Way ANOVA |
| ISI | 2 | F = 31.689 | **P < 0.001** | Two-Way ANOVA |
| Group x ISI | 3 | F = 2.171 | P = 0.121 | Two-Way ANOVA |
| **SCI CPN-TN vs TN-TN** | | | | |
| Group | 1 | F = 6.503 | **P < 0.012** | Two-Way ANOVA |
| ISI | 2 | F = 1.894 | P = 0.155 | Two-Way ANOVA |
| Group x ISI | 3 | F = 0.668 | P = 0.514 | Two-Way ANOVA |

**Supplemental Table 5.** Fig. 9 Results of one and two-way ANOVAs comparing % change of H-reflex across long ISIs (0.5 to 2 s) and between SCI and control groups for the 1.0 x MT and 1.5 x MT conditioning CPN stimulation intensities.

| Figure 10: H-reflexes in SCI participants not taking and taking baclofen and controls | | | | |
|---|---|---|---|---|
| **Fig. 10A: TN-TN** | **SCI no bacl.** | **SCI bacl.** | **Control** | |
| % chg H-reflex | -46.71 (4.67) | -65.18 (4.62) | -70.32 (0.70) | |
| SCI no bacl vs SCI bacl | t = 6.286 | **p < 0.001** | Student's t test | |
| SCI no bacl vs control | t = 2.459 | **p = 0.039** | Student's t test | |
| **Fig. 10B: CPN-TN** | **DF** | **Test value** | **P value** | **Test** |
| Group | 2 | F = 35.023 | **P < 0.001** | Two-way ANOVA |
| ISI | 4 | F = 8.391 | **P < 0.001** | Two-way ANOVA |
| Group x ISI | 8 | F = 2.388 | **P = 0.020** | Two-way ANOVA |
| **0.5 s ISI** | | | | |
| SCI no bacl vs Control | | t = 7.148 | **P < 0.001** | Holm-Sidak |
| SCI no bacl vs SCI bacl | | t = 3.395 | **P < 0.001** | Holm-Sidak |
| SCI bacl vs Control | | t = 3.227 | **P = 0.002** | Holm-Sidak |
| **1.0 s ISI** | | | | |
| SCI no bacl vs Control | | t = 3.320 | **P = 0.001** | Holm-Sidak |
| **1.5 s ISI** | | | | |
| SCI no bacl vs Control | | t = 3.496 | **P < 0.001** | Holm-Sidak |
| SCI no bacl vs SCI bacl | | t = 2.869 | **P = 0.005** | Holm-Sidak |
| **2.0 s ISI** | | | | |
| SCI no bacl vs Control | | t = 2.819 | **P = 0.006** | Holm-Sidak |
| SCI no bacl vs SCI bacl | | t = 2.578 | **P = 0.011** | Holm-Sidak |

**Supplemental Table 6. A)** Mean and SD of last 5 H-reflexes during a 1 Hz stimulus train for participants with SCI taking and not taking oral baclofen and controls and results of unpaired comparisons between groups. **B)** Results of two-way ANOVAs comparing % change of H-reflex across the long ISIs (0.5 to 2.5 s) and between the three groups.
